# Chemostat culturing reduces fecal eukaryotic virus load and delays diarrhea after virome transplantation

**DOI:** 10.1101/2024.11.20.624498

**Authors:** Simone Margaard Offersen, Signe Adamberg, Malene Roed Spiegelhauer, Xiaotian Mao, Torben Sølbeck Rasmussen, Frej Larsen, Jingren Zhong, Duc Ninh Nguyen, Dennis Sandris Nielsen, Lise Aunsholt, Thomas Thymann, Kaarel Adamberg, Anders Brunse

## Abstract

Fecal virome transfer (FVT) shows promise in reducing necrotizing enterocolitis (NEC), likely due to donor bacteriophages preventing the gut dysbiosis preceding disease. However, concurrent transfer of eukaryotic viruses may carry a risk of infection for the recipient. To increase safety, we investigated chemostat propagation as a method to eliminate eukaryotic viruses from donor feces while maintaining a diverse and reproducible bacteriophage community. Donor feces was collected from healthy suckling piglets and inoculated into a fermenter containing growth media supplemented with lactose and milk oligosaccharides (MOs). During continuous medium exchange (20% volume/h), dilution significantly reduced eukaryotic viruses. Viral richness was concurrently reduced although still preserving a stable community of 200-250 bacteriophages. Inclusion of MOs in the medium ensured higher bacterial richness and a bacterial community closer resembling donor feces. Fecal *Lactobacillaceae* bacteria were lost during cultivation but partially replaced by members of the *Bacteroidota* phylum in MO-supplemented cultures, accompanied by phages predicted to have *Parabacteroides* as host. After cultivation, virus-like particles (VLPs) were isolated, and their ability to reduce NEC incidence tested *in vivo*. Preterm piglets were delivered by cesarean section and received either the lactose- or MO-propagated viromes by oral route (*n* = 14-15/group). These were compared with groups receiving the same dose of donor fecal virome (10^10^ VLPs/kg) or vehicle control. The piglets were subsequently fed infant formula for 96 hours followed by euthanasia and tissue sampling. Both chemostat-propagated viromes effectively mitigated diarrhea compared to the donor virome. The donor virome partially engrafted in recipients and led to higher levels of *Lactobacillaceae* bacteria and *Lactobacillaceae* targeting phages. However, these signatures were lost in recipients of chemostat-propagated viromes, and only minor microbiome effects and no NEC prevention were observed. To conclude, we provide *in vivo* proof-of-concept for chemostat propagation of fecal viruses as a means to deplete eukaryotic viruses and in turn reduce side effects in newborn virome recipients. However, chemostat culture conditions need further optimization to preserve the donor phageome.

## Introduction

Immediately after birth, microorganisms colonize the gastrointestinal tract, thereby establishing the gut microbiome (GM). Non-digestible milk oligosaccharides (MOs) present in breast milk strongly influence GM assembly by acting as highly specific microbial substrates to promote the growth of bifidobacteria and *Bacteroides*^1^. Preterm birth (< 37 weeks of gestation) is often accompanied by GM perturbations due to factors such as cesarean section, early antibiotic use, immature gastrointestinal tract, and partial formula feeding^2–5^. These disturbances inhibit beneficial anaerobes like bifidobacteria and *Bacteroides* whereby facultative pathobionts like *Enterobacteriaceae* members may take over and initiate disease processes^6^. This can ultimately lead to the severe intestinal disorder necrotizing enterocolitis (NEC), characterized by inflammation and necrosis of intestinal tissue^7^. With global mortality rates ranging from 20-50%, NEC is considered a critical medical and surgical emergency^8^.

In recent years, there has been growing interest in studying the gut virome alongside gut bacteria, particularly in the context of severe inflammatory gut disorders like NEC and inflammatory bowel disease (IBD). Specifically, bacteriophages (phages for short) are abundant gut viruses that function to modulate bacterial growth through host-specific lysis or lysogeny^9^. This dynamic interaction and additional preference for intestinal mucosal sites enable phages to alter bacterial compositions and protect the host from bacterial invasion^10–12^. Several studies have associated virome alterations with IBD^13–15^ and recently, the link was discovered in a small cohort of NEC patients as well^16^. Thus, the virome may be a novel target for modulating the early life GM and preventing intestinal inflammation in neonates.

Fecal virome transfer (FVT) shows potential for restoring gut health by introducing viruses from donor feces to patients of gut dysbiosis. A recent preclinical study demonstrates that FVT can modify the gut virome of preterm piglets, reducing mucosa-associated *Enterobacteriaceae* levels and providing significant protection against NEC^17^. However, during FVT, fecal eukaryotic viruses are transferred alongside phages, introducing a risk of infections. While screening programs exclude fecal donors with known viral pathogens, many eukaryotic viruses in the gut have unknown functions that could affect neonatal health, potentially leading to both immediate and long-term consequences^18,19^. A follow-up study using the piglet model found that, while a prescreened FVT indeed offered protection against NEC, it was also linked to adverse events, namely an earlier onset of diarrhea, which also led to significant weight loss^20^.

Fecal phages and eukaryotic viruses are highly similar in size and structure^21^. As a result, phages will be damaged if traditional methods for eliminating eukaryotic viruses, such as chemicals or irradiation, are applied to FVT preparations. However, both types of viruses rely on host cells for propagation but have very distinct host specificities (bacterial vs. mammalian cells). This distinction can be leveraged in a *chemostat*: a bioreactor system designed to cultivate microorganisms in a controlled environment^22–24^. Here, a continuous fecal culture is maintained by continually adding fresh medium, providing nutrients for bacterial growth, and thus continuously generating hosts for phage propagation. Concurrently, equivalent volumes of spent media containing microorganisms and metabolic by-products are removed, leading to a gradual dilution of eukaryotic viruses, whose host cells are unable to propagate in the chemostat^24,25^. Thus, the method has the potential to create a steady-state fecal culture from which the phage communities can be extracted and used for FVT^26,27^.

Insights from traditional fecal microbiota transplantations (FMT) for IBD in humans or experimental NEC suggest that treatment efficacy can vary depending on the donor, a phenomenon that likely applies to FVT as well^28–30^. In the previous NEC studies, feces from suckling term-born pigs were used as FVT donor material to best match the preterm piglet recipients^17,20^. Although this donor-recipient matching approach seems promising, the use of young donors limits the amount of fecal matter available for donation. The chemostat method overcomes these limitations by generating a mass-reproducible fecal phage community, potentially providing both a safer and donor-independent FVT strategy.

To propagate bacteria and phages from young donors, the medium composition should replicate the nutrients reaching the gut during nursing. Thus, we tested if adding medium MOs would increase the microbial similarity between the donor feces and chemostat culture. Subsequently, native fecal virome transfer (FVT) was compared with the chemostat virome transfer propagated without MOs (CVT) and with MOs (CVT-MO) in the preterm piglet model. We hypothesized that chemostat-propagated viromes could modulate the GM and reduce NEC lesions while avoiding potential side effects, such as earlier onset of diarrhea.

## Materials and methods

### Donor feces collection

Gut content was collected from four 10-day-old healthy-appearing piglets from four different conventional farms. After euthanasia, the cecum, colon, and rectum were isolated, and the content was emptied and processed under anaerobic conditions. After diluting 1:1 in 2 × SM buffer (400 mM NaCl, 20 mM MgSO4, 100 mM Tris-HCl, pH 7.5) containing 20% sterile glycerol, the content from each pig was homogenized, divided into smaller fecal aliquots, and stored at −70°C.

### Chemostat cultivation Growth medium

A rich tryptone and yeast extract based medium supplemented with mineral salts and vitamins was prepared as described by Pham et al.^23^ with the following modifications: lactalbumin concentration 6 g/L and 0.05 M potassium phosphate buffer were used. Additionally, carbohydrate content varied as follows: In one setup, a mixture of MOs was added (7 g/L, LAC-MO medium) along with the original content of lactose (6.4 g/L) and mucin (4 g/L porcine gastric type II mucin, Sigma-Aldrich, Germany). The total MO level was adjusted to quantities reported in mature porcine milk^31^. This consisted of 3.5 g/L 3’-sialyllactose (3’SL), 2.8 g/L lacto-N-neotetraose (LNnT), and 0.7 g/L 2’-fucosyllactose (2’FL), chosen to match porcine milk levels of sialylated-, neutral-, and fucosylated MOs, respectively^32^. All MOs were donated by DSM-Firmenich (Switzerland). In another setup, the MO fraction was substituted with lactose, balanced by equimolar levels of C6 monosaccharides. This resulted in 12.5 g/L lactose (LAC medium) in addition to the 4 g/L mucin.

### Cultivation system and culture conditions

The previously described chemostat system^22,24^ was used for cultivation of the collected pig feces. Briefly, the cultivation system consisted of an anaerobic fermenter equipped with sensors for pH, pO2, and temperature control. The culture volume (V) was kept constant (800 mL) as well as the temperature (39°C)^33^ and pH (6.5). The pH was controlled by adding 3 M NaOH according to the setpoint. Variable speed pumps enabled a controlled inflow and outflow of medium during continuous cultivation. The collected fecal aliquots were thawed and processed under anaerobic conditions where all four donors were pooled in equal amounts and further diluted 1:1 in sterile 0.9% saline (resulting in four times diluted feces). To start the experiments, 4 mL of diluted fecal pool was inoculated into the fermenter (0.125% w/v raw feces in fermenter volume) and allowed growth for 24 hours in a fixed medium volume, which corresponds to the end of the exponential growth phase of the fecal culture (batch phase). Hereafter, continuous medium inflow/outflow was started (chemostat phase) at a feeding rate (F) of 160 mL per h, resulting in a dilution rate (D) of 0.2 h^-1^ (D=F/V). This phase continued until 75 h, during which the chemostat volume was exchanged 10 times (Fig. 1A). Two and four replicates were carried out with LAC and LAC-MO medium, respectively. Samples from the outflow were collected on ice three times during the batch phase and four times during the chemostat phase as shown in Fig. 1A. On-line and at-line parameters used for experiment control are depicted in Supplementary Fig. S1.

**Figure 1.**
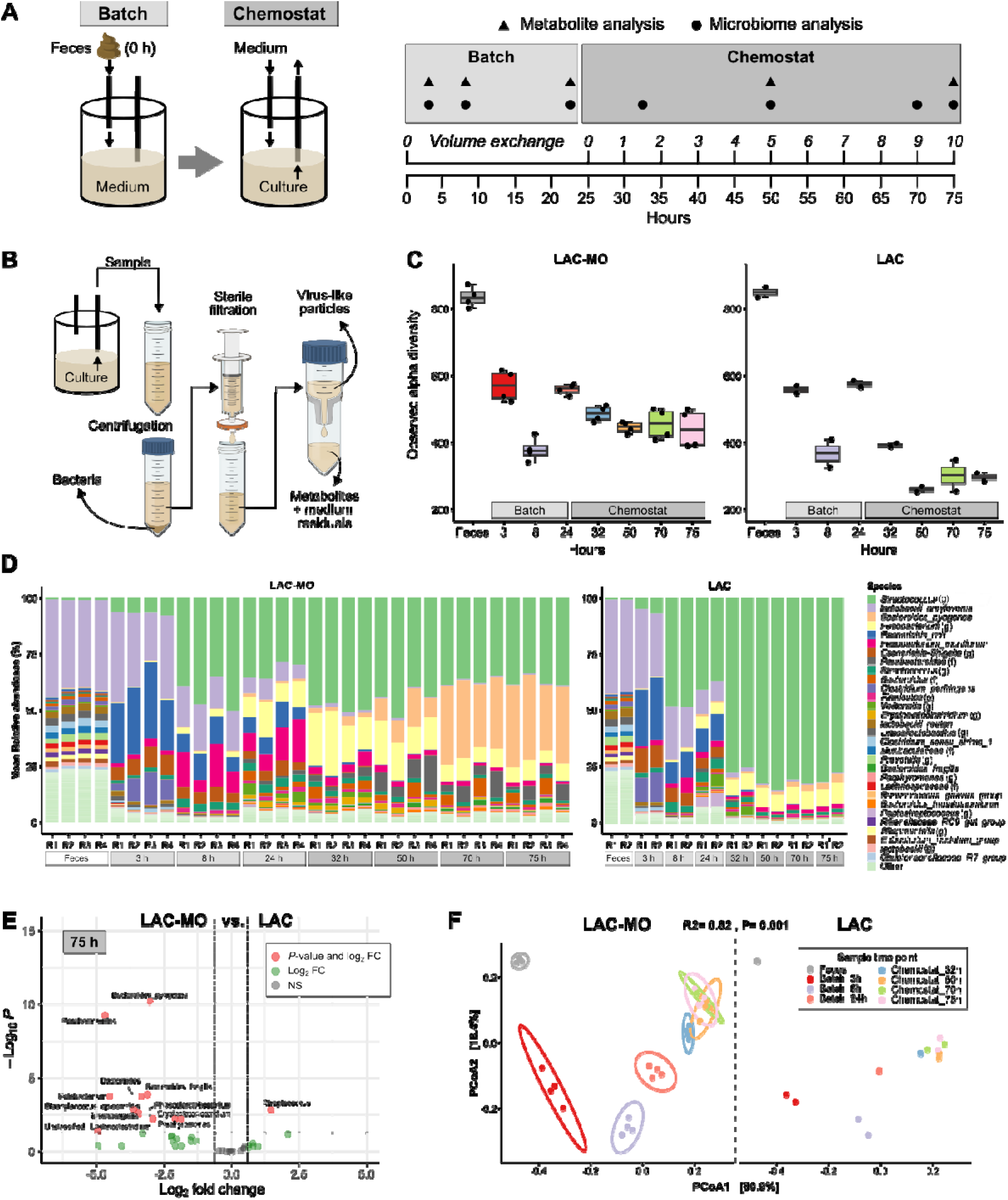
MO supplementation preserves bacterial richness and promotes *Bacteroidota* enrichment in stabilized chemostat cultures. A) The experimental setup of chemostat cultivations of donor feces with lactose medium (LAC, *n*=2) or lactose and milk oligosaccharide medium (LAC-MO, *n*=4). Donor feces was inoculated into a fermenter followed by batch growth for 24 h. The chemostat culture was then run until 10 volume exchanges. The sampling points for metabolite and microbiome analyses are indicated. B) Processing of chemostat culture to separate virus-like particles from bacteria and the culture residuals. C) Number of observed bacterial species. D) Relative bacterial abundance per replicate summarized at species level. Taxonomic rank specifications: p = phylum, f = family, g = genus. E) Volcano plot visualizing bacteria DESeq2 enrichment analysis with FDR correction based on 16S rRNA gene amplicon sequencing on species level. Horizontal dotted lines show a significance level of *P* < 0.05 and vertical lines show ± 0.6 log2 fold change (FC) cut-off. NS = non-significant. F) Principal coordinate analysis (PCoA) plot visualizing bacterial beta diversity over time, based on Bray-Curtis dissimilarity metrics.

### Gas, redox, biomass, and chromatographic analyses

The composition of the gas outflow, including H, CO, HS, CH, and N, was evaluated using an Agilent 490 Micro GC Biogas Analyzer (Agilent 269 Technologies Ltd., USA) connected to a thermal conductivity detector. Gas flow volume was monitored regularly using a MilliGascounter (RITTER Apparatebau GMBH & Co, Germany). The redox potential of both the growth medium and culture supernatant was measured with a pH/Redox meter and an InLab® Redox electrode (Mettler Toledo, USA). Biomass dry weight was assessed gravimetrically from a 10 mL culture by centrifuging at 6,000 rpm for 20 minutes, followed by washing the biomass with distilled water and drying it at 105°C for 20 h in an oven.^24^

### Chromatographic analyses

Culture samples were centrifuged (11,000 × *g* for 5 min at 4°C), then stored at −20°C. After thawing, the supernatants were filtered using Amicon® Ultra-10K Centrifugal Filter Devices with a 3 kDa cut-off, following the manufacturer’s instructions (Millipore, USA). Organic acids (succinate, lactate, formate, acetate, propionate, isobutyrate, butyrate, isovalerate, and valerate) and ethanol concentrations were measured via high-performance liquid chromatography (HPLC) using the Waters Alliance 2795 system (Waters, Milford, MA, USA) with a BioRad HPX-87H column (Hercules, CA, USA). Isocratic elution with 0.005 M HSO was performed at a flow rate of 0.5 mL/min at 35°C, with refractive index (RI) (model 2414) and UV (210 nm; model 2487) detectors (Waters, USA). Quantification was done using analytical-grade standards, with a detection limit of 0.1 mM.^24^

### Pre-processing of samples for separation of bacterial and viral analysis

Fecal inoculum and culture samples were analyzed to investigate microbiome changes over time. Separation of the viruses and bacteria was done as illustrated in Fig. 1B. Fecal inoculum was diluted in SM buffer (200 mM NaCl, 10 mM MgSO4, 50 mM Tris-HCl, pH 7.5) until 1:20 and centrifuged (4,500 × *g,* 30 min, 4°C). For chemostats, 10 mL cooled outflow culture were sampled per time point and immediately centrifuged (4,500 × *g,* 30 min, 4°C). The pellet was stored at −80°C until bacterial analysis and the supernatant was filtered through a 0.45 µm Filtropur S PES syringe filter (Sarstedt, Helsingborg, Sweden). Subsequently, the filtrate was concentrated using Vivaspin 20 units with a 300 kDa molecular weight cut-off (MWCO) PES membrane (Sartorius, Göettingen, Germany), which was centrifuged (2,500 × *g,* 4°C) until all filtrate volume had passed. Hereafter, 2 mL SM buffer was added and stored at 4°C overnight to release viruses from the membrane. The concentrated virus samples were stored at −80°C until analysis.

### Processing of chemostat culture and feces for separation of viruses

After 10 volume exchanges, the total fermenter culture (800 mL) was drained into a sterile 1,000 mL bottle on ice. Thereafter, the culture was centrifuged (7,000 × *g,* 60 min, 4°C) and the supernatant was filtered to remove bacteria using 0.45 µm Filtropour V50 PES vacuum filtration units (Sarstedt, Helsingborg, Sweden). To concentrate viruses and remove residual medium components, including MOs, the filtrate was ultracentrifuged following a protocol shown to retain infective viruses^34^. To further reduce non-viral particle contamination, we employed a higher MWCO of 300 kDa, compared to the original protocol. Thus, the filtrates were added to Vivaspin 20 units (300 kDa MWCO, PES; Sartorius, Göettingen, Germany) and centrifuged (6,000 × *g* at 25° fixed angle, 4°C). Each unit had 80 mL repeatedly added and concentrated to 0.5-1.0 mL. Hereafter, SM buffer was added to 10 mL total and stored at 4°C overnight. The eight times concentrated culture virome from all units were pooled, then aliquoted and stored at −70°C until the piglet experiment. This procedure was repeated for each chemostat replicate.

Fecal virome transfer (FVT) solution was prepared similarly by pooling the four donors and further diluting 1:5 in SM buffer. The fecal viromes were isolated as described above (Fig. 1B). After concentration with Vivaspin 20 units, SM buffer was added to the original volume (1:10 diluted). Aliquots of FVT and sterile SM buffer were stored at −80°C until use.

### Fluorescence microscopy and bacterial cultivation

Epifluorescence microscopy was used to determine the concentration of virus-like particles (VLPs) in each of the end culture viromes and the FVT. VLPs were stained with SYBR Gold (Cat. no S11494, Invitrogen, ThermoFisher Scientific) and captured on 0.02 µm aluminum oxide filters (Whatman™ 6809-6002 Anodisc™, Cytiva, USA). One filter was prepared for viromes from each chemostat run and the fecal inoculum. The filters were photographed (Photometrics CoolSNAP camera) using an epifluorescence microscope fitted with a 490 nm filter block. Hereafter, VLPs were counted and the number per mL was estimated as described previously^20^. To confirm the absence of living bacteria in each virome solution, 100 uL from aliquots of CVT, CVT-MO, FVT, and SM buffer (control, CON) was spread on plate count agar. The plates were incubated for 24 hours under both aerobic and anaerobic conditions for 24 h before inspection for bacterial growth. A plate with no added aliquots and a plate with 100 uL of fecal slurry were included as negative and positive control, respectively.

### Cell stimulation

The human monocyte cell line THP-1 was used to investigate the innate immune response following challenge with the fecal and chemostat viromes compared with a cocktail of 10 isolated virulent phages, and an *Escherichia coli* bacterium. A fecal virome transfer (FVT) aliquot was thawed and used directly. Chemostat virome transfer (CVT) solution consisted of pooled viromes from both chemostats propagated with LAC medium (replicas 1 and 2). Similarly, chemostat virome transfer propagated with MOs (CVT-MO) comprised pooled viromes from two of the four chemostats propagated with LAC-MO medium (replicas 1 and 3). Each fecal or chemostat solution (FVT, CVT, CVT-MO) was ultracentrifuged (300 kDa MWCO) and adjusted to 5 × 10^9^ VLPs per mL. The phage solutions were mixed to reach a concentration of 5 × 10^9^ plaque-forming units (PFU) per mL, as described by Larsen et al.^34^. Details of the phages are listed in Supplementary Table S1. The *E. coli* strain (ST-2064) was cultured in a liquid Luria-Bertani medium (BD Difco™ LB Broth Lennox, Becton Dickinson, USA) until it reached the exponential phase and adjusted to a concentration of 10^8^ colony forming units (CFU) per mL. Before the challenge, endotoxins were removed from all virome, phage, and bacterial preparations using Pierce™ High Capacity Endotoxin Removal Spin Columns (0.5 mL, ThermoFisher Scientific, USA) by following the manufacturer’s instructions. The THP-1 cells were cultivated, stimulated, and RNA was extracted and RT-qPCR performed exactly as described previously^20^. In brief, THP-1 cells were differentiated into macrophages and stimulated with 100 µL of FVT, CVT, CVT-MO, phage solution (Phages), or *E. coli* solution. RNA was extracted after incubation for 20 h (37°C, 5% CO_2_). Seeding and challenges were performed in triplicates at cell passages 17, 19, and 20.

### Piglet Experiment 1

Experimental procedures conducted during the animal experiments were approved by the Danish Animal Experiments Inspectorate (license number: 2020-15-0201-0052), under the guidelines from Directive 2010/63/EU of the European Parliament. 60 piglets were delivered preterm from three sows by cesarean section (day 106, 90% gestation) as described previously ^35^. After delivery, the pigs were transferred to heated incubators and observed or ventilated until the respiration stabilized. Hereafter, the pigs were fitted with oral catheters for enteral feeding and umbilical arterial catheters for parenteral nutrition (PN) and access to blood circulation (6 and 4 Fr size, respectively, Portex, Kent, UK). The pigs were stratified by sex and birth weight and randomly allocated to four groups receiving fecal virome transfer (FVT), chemostat virome propagated with LAC medium (CVT), chemostat virome propagated with LACMO medium (CVT-MO), or SM buffer (control, CON).

### Immunization and feeding procedures

To substitute colostrum feeding, all pigs were immunized with sows’ plasma (16 mL/kg) by intraarterial infusion over two hours on the first day of life. Hereafter, the pigs received declining levels of PN over the study period (96-48 mL/kg^0.75^/d, Kabiven, Vamin, Fresenius-Kabi; Bad Homburg, Germany). Enteral nutrition (EN) was fed every three hours and consisted of a home-mixed formula (formula 1) as described previously^20^. EN volumes increased gradually, reaching 24, 40, 64, and 96 mL/kg^0.75^/d on days 1, 2, 3, and 4– 5, respectively.

### Fecal- and chemostat virome transfers

CVT and CVT-MO were produced as described for the cell stimulation assay above. Just before use, the treatments were thawed and then administered orally via the orogastric tube. Each virome treatment (FVT, CVT, CVT-MO) was adjusted to 10^10^ VLP/kg in a total volume of 2-3 mL/kg SM buffer. The CON group received 2 mL/kg of SM buffer. All groups received their treatment volume divided into administrations twice daily on days 1 and 2 (Fig. 4A).

### In vivo procedures and euthanasia

The pigs were closely monitored by experienced personnel and pigs showing clinical signs of NEC (abdominal distention, lethargy, or severe bloody diarrhea) were immediately euthanized. Daily weights and exact times of diarrhea episodes were recorded. On days 3 and 5, blood samples were obtained for routine hematological profiling. On day 5, all pigs were anesthesized followed by euthanasia with an intracardiac injection of pentobarbital. As a marker for intestinal permeability, the urinary lactulose-to-mannitol ratio was measured as previously described^36^.

### NEC evaluation and tissue sampling

The abdominal organs were excised, separated, and weighed. The gastrointestinal tract was assessed for NEC-like lesions as previously described^37^. Briefly, levels of hyperemia, hemorrhage, pneumatosis intestinalis (intramural gas), and necrosis were assessed by a blinded investigator and converted to a 16-point NEC score. NEC was defined as the presence of hemorrhage, pneumatosis intestinalis, and/or necrosis. Colon content, two samples from the proximal intestine, as well as mesenteric lymph nodes from the ileocecal region were snap-frozen for later analysis. The most severe lesions from the small intestine and colon were both snap-frozen and fixed in 4% paraformaldehyde. The fixed tissue was later paraffin-embedded, sectioned, and stained with hematoxylin and eosin. A blinded pathologist assessed each section by microscopy and assigned an established histopathological score ranging from normal appearance (score 0) to transmural necrosis (score 8)^37^. Lactase activity was measured in proximal small intestinal tissue by a previously described colorimetric method^36^. Interleukin (IL) 1β and IL-8 were measured in tissue homogenates from proximal small intestine and colon lesions by commercial porcine ELISA kits (R&D Systems, 158 Abingdon, Oxfordshire, United Kingdom) according to the manufacturer’s instructions^38^.

### RNA sequencing of mesenteric lymph nodes

Mesenteric lymph node RNA was extracted from 20 mg homogenized tissue (GentleMACS Dissociator, Miltenyi Biotec, Bergisch Gladbach, Germany) using the RNeasy Mini Kit (Qiagen, Hilden, Germany) following the protocol for tissue, and with on-column DNase digestion (RNase-free DNase, Qiagen, Hilden, Germany). Whole-transcriptome shotgun sequencing was performed by Novogene services (Cambridge, UK) and analyzed as described previously^39^. Briefly, RNA-Seq libraries were constructed using Novogene NGS RNA Library Prep Set (PT042) and sequenced using the Illumina NovaSeq 6000 S4 flowcell platform. Fastp software was utilized for trimming the raw reads to remove adapters and low-quality bases. The raw sequencing data were saved in FASTQ format, containing both the read sequences and their associated base quality scores. HISAT2 was employed to align the reads to the porcine reference genome (*Sscrofa11.1*). FeatureCounts package was then used to create a gene count matrix, which was further analyzed for differentially expressed genes (DEGs) with DESeq2, using *P*-value cutoff 0.05 and log2 fold change cutoff 1. The pathway enrichment analysis was performed with a gene set enrichment analysis (GSEA) through the clusterProfiler package using the Gene Ontology (GO) classification system for *Sus scrofa*. The false-discovery-rate (FDR) cut-off of 0.05 was applied to identify significant pathways. A normalized enrichment score (NES) was obtained, with either negative or positive values indicating the pathways being down- or up-regulated. The enriched pathways were visualized using ggplot2.

### Piglet Experiment 2

53 piglets delivered from two sows were stratified by sex and birth weight and randomly allocated to two groups receiving CVT-MO solution or SM buffer (CON). The enteral nutrition (EN) consisted of an ultra-high temperature (UHT) treated ready-to-drink formula (formula 2), previously reported to have a high capacity for inducing NEC lesions ^37^. All other experimental procedures were exactly as described for *Piglet Experiment 1*.

### Bacterial DNA extraction, sequencing, and pre-processing of raw data

The DNAeasy PowerSoil Pro Kit (Qiagen) was used to extract bacterial DNA from the fecal bacterial pellet and the chemostat aliquot following the manufacturer’s instructions. The final purified DNA concentration was determined on Varioskan Flash (Thermo Fisher Scientific, USA) using Qubit™ 1x dsDNA high sensitivity kit (Invitrogen). The purified DNA products were stored at −80°C for later use. The full-length 16S rRNA gene was used to determine the gut bacterial composition. The 16S rRNA gene was amplified using several forward and reverse primers (Suppl. Table S2). The first PCR reaction included mixing of 12 μL PCRBIO Ultra Mix (PCRBIO), 6 μL Sterile Milli Q H2O, 2 μL primer mix, and 5 μL genomic DNA (∼5 ng/μL). The PCR cycling was run on SureCycler 8800 (Agilent Technologies). The first PCR cycles were as follows: 95°C for 2min; 2 cycles of 95°C for 20sec, 48°C for 30 sec, 65°C for 10sec, and 72°C for 45 sec; followed by a final step at 72°C for 4min. The second PCR step was done by barcoding the 11 μL of the first PCR products with 2 μL barcodes in 12 μL PCRBIO Ultra Mix (PCRBIO): 95°C for 2min; 33 cycles of 95°C for 20 sec, 55°C for 30 sec, and 72°C for 40 sec; followed by a final step at 72°C for 4min. The gut bacterial composition was assessed using the Oxford Nanopore Technologies (ONT) GridION platform. ONT MinKNOW software v22.10.7 collected the data generated by GridION. The Guppy v6.2.8 base-calling toolkit was used to convert the the raw FAST4 files to FASTQ format. The Long Amplicon Consensus Analysis (LACA) pipeline generated the abundance table through raw FASTQ files (https://github.com/yanhui09/laca). The assigned taxonomy quality was corrected using SILVA database^40^.

### Viral RNA/DNA extraction, sequencing, and pre-processing of raw data

The viral particle separation was conducted based on an existing protocol^41^. The intestinal content from pig and chemostat aliquots was suspended in 5 mL autoclaved SM buffer (100mM NaCl, 8mM MgSO4·7H2O, 50mM Tris-HCl with pH 7.5) followed by centrifuging at 4500 x *g* for 30 min at 4 °C. The supernatant was filtered using a 0.45µm Minisart High Flow PES syringe filter (Sartorius) to remove bacteria and other larger particles. VLPs in the filtered supernatants were concentrated and smaller metabolites removed by ultrafiltration using a Centrisart ultrafiltration devices with a filter cutoff at 100 kDa (Sartorius, UK). To remove free DNA/RNA molecules, 140 µL supernatants were treated with 5 units of Pierce Universal Nuclease (ThermoFisher Scientific) for 15 min at room temperature. Viral DNA/RNA was extracted with Viral RNA mini kit (Qiagen). Reverse transcription was conducted using SuperScript VILO Master (Thermo Fisher Scientific) and subsequently cleaned with DNeasy blood and tissue kit (Qiagen) by following steps 3-8 in the manufacturer’s manual. Multiple displacement amplification was performed to include ssDNA viruses using the GenomoPhi V3 DNA amplification kit (Cytiva). The library preparation for sequencing was conducted by using Nextera XT kit (Illumina) as previously described^26^. The pooled purified PCR product was sequenced on a NovaSeq X plus platform by Novagene. Raw reads were trimmed (trimmomatic v0.38), deduplicated (seqkit v0.13.2) and assembled (spades v3.13.0)^42–44^. Assembled contigs were deduplicated at species level (∼95% sequence identity) and non-viral contigs were removed by running geNomad^45^ and checkV^46^ and selecting contigs that were either 1) marked “High-quality” or “Complete” by checkV, or 2) had a geNomad score of >0.9 and >2 hallmark genes. Additionally, all viral contigs should be above 2200 bp, or 10.000 bp if the geNomad-predicted taxonomies were caudoviral. Viral hosts were predicted using iPHoP^47^ and taxonomies were set by aligning to a custom database. Protein prediction was done with phold (https://github.com/gbouras13/phold), which was used to predict whether a phage was temperate or not based on the presence of recombinase/integrase genes. Reads from each sample were mapped to the viral contigs by msamtools (v1.1.2). The pipeline is described in detail at https://github.com/frejlarsen/vapline3.

### Bioinformatic analysis of bacterial and viral sequences

R version 4.3.0 was used for analysis and presentation of microbiome data. The main packages used were phyloseq^48^, vegan^49^, DESeq2^50^, ampvis2^51^, ggpubr, psych, EnhancedVolcano, and ggplot^48^. The R package microDecon (runs=1, regressions=1) was used to remove the contamination of viral contigs by read counts in negative controls^35^, and 83.0% were kept after removal. For samples in the chemostat experiment, the media used in the chemostat propagation was used as a control to remove the viral contigs introduced by the media, using microDecon^52^ (runs=1, regressions=1), leaving 84.2% contigs for analysis of chemostat samples. Cumulative sum scaling normalization was performed using R metagenomicSeq package. The Bray-Curtis dissimilarity metric was used for beta diversity analysis and with statistics based on one-sided pairwise PERMANOVA corrected by FDR. The differential bacteria and viruses were identified by DESeq2 on the summarized bacterial species level and viral contigs (vOTUs) level based on two-sided Wald test. Alpha diversity and shared vOTUs were analyzed with the non-parametric two-sided Wilcoxon rank-sum tests with FDR correction. Eukaryotic viral contigs were manually identified in the vOTU table and the relative abundance of these contigs was calculated based on total eukaryotic viral taxa and subsequently visualized in heatmaps.

### Statistical analysis of remaining data

Remaining statistical analyses were performed in R version 4.2.2. Survival and diarrhea onset were analyzed with pairwise Log-Rank tests and Benjamini-Hochberg (BH) adjustment. Continuous outcomes were analyzed with linear mixed models with group, sex, and birth weight as fixed effects and litter as a random effect. The group effect was assessed with an ANOVA, followed by Tukey’s test. To ensure the validity of the modeled data, the residuals and fitted values were evaluated for normality and homogeneity of variance. For some outcomes, this necessitated log-transforming before modeling. Tissue cytokines did not meet parametric criteria and were analyzed with the Kruskall-Wallis test followed by Dunn’s test with BH correction. Gross pathology and microscopic lesion scores were analyzed using a proportional ordered logistic regression model (R package MASS). The analysis included group, sex, litter, and birth weight as fixed effects. Group effects were assessed using the likelihood ratio test with a BH correction. Incidences of pathologies were compared with Fisher’s exact tests. A repeated measures linear mixed model was employed to analyze chemostat metabolite levels, with fixed effects for group, time, and their interaction, and a random intercept for replicate ID. Piglet repeated measures were analyzed using a linear mixed model, with fixed effects for group, day, their interaction, and covariates (sex, litter, birth weight). To account for the correlation and variance heterogeneity over time, this model assumed an unstructured covariance pattern (R package LMMstar).

## Results

With this study, we aimed to provide a combined *in vitro* and *in vivo* proof-of-concept for phage preservation and concomitant eukaryotic virus reduction through chemostat propagation of fecal matter and subsequent virome transfer into newborn recipients to alleviate NEC and virus-associated diarrhea. Further, we measured the effect of supplementing the chemostat cultures with MOs for better approximation of a milk-shaped fecal inoculum.

### Milk oligosaccharides preserved the bacterial richness of a stable fecal chemostat culture

A mix of anaerobically collected fecal samples from four healthy suckling pigs was inoculated into batch cultures with or without an MO blend representing the major MOs in porcine milk to supplement lactose as the major carbohydrate source^32,53,54^. Continuous exchange of medium was initiated after 24 hours of incubation and a constant target pH of 6.5 (mean 6.55 ± 0.04 at 50-75 hours for all chemostats) was maintained by base titration. To check the stability of the chemostat cultures, base titration curves were inspected (Suppl. Fig. S2). In the batch phase, lactose conversion to lactic acid resulted in a peak titration at 4-5 hours with a higher peak in the pure lactose (LAC) cultures. A later pattern of small but prolonged titration occurred only in the MO (LAC-MO) supplemented cultures, indicating a slower utilization of MOs. After initiation of continuous volume exchange, titration patterns quickly stabilized with marginally higher acid production/base consumption in the LAC cultures.

We assessed the bacterial composition of the cultures at seven time points throughout the experiment to assess culture dynamics (Fig. 1A-B). Whereas all cultures lost a significant proportion of bacterial richness already in the early batch phase, supplementing the growth medium with MOs preserved the richness of the culture throughout the following chemostat phase to a greater extent than pure lactose (Fig. 1C). Taxonomically, donor feces showed >50% relative abundance of bacteria from the *Lactobacillaceae* family. During the early batch phase, *Escherichia coli* expanded rapidly, contributing to the fast lactose consumption and acid production. Hereafter, both *Lactobacillaceae* and *E. coli* were replaced mainly by *Streptococcus* and *Fusobacterium* spp. Importantly, in the chemostat phase a significant proportion of these were replaced by different species of the *Bacteroidota* phylum only in the MO cultures indicating that these taxa had a competitive advantage due to MO utilization capabilities (Fig. 1D). Consequently, in the MO-supplemented end culture we found a relative enrichment of several taxa, all of which were obligate anaerobes (Fig. 1E).

Compositional analysis using Bray-Curtis ordination revealed an abrupt change in the early batch phase, with a partial restoration towards the composition of the fecal inoculum occurring throughout the chemostat phase visualized along the second principal coordinate. Notably, MO supplementation augmented this restoration. After five volume exchanges, the bacterial composition had stabilized and did not develop further for the remainder of the experiment. However, using the current fermentation conditions including a relatively high dilution rate, the chemostat end culture did not fully approximate the composition of the fecal inoculum (Fig. 1F).

Metabolically, the culture dynamics throughout the experiment corroborated the bacterial compositional analysis. As such, high production of lactate, formate, ethanol, succinate, H_2,_ and CO_2_ in the early batch phase indicated mixed-acid fermentation mainly by *E. coli* (Fig. 2). As the cultures entered the chemostat phase, lactate was gradually consumed, and short-chain fatty acid levels (e.g. propionate, butyrate) increased. In the stable end cultures, butyrate levels were equally high, but importantly, propionate was a major fermentation product only in the MO cultures. Other short-chain fatty acids (e.g. valerate, isovalerate) were also increased by MOs despite lower concentrations. Taken together, the increased proportion of obligate anaerobes in the MO-supplemented chemostat cultures greatly increased the production of propionate and other short-chain fatty acids.

**Figure 2.**
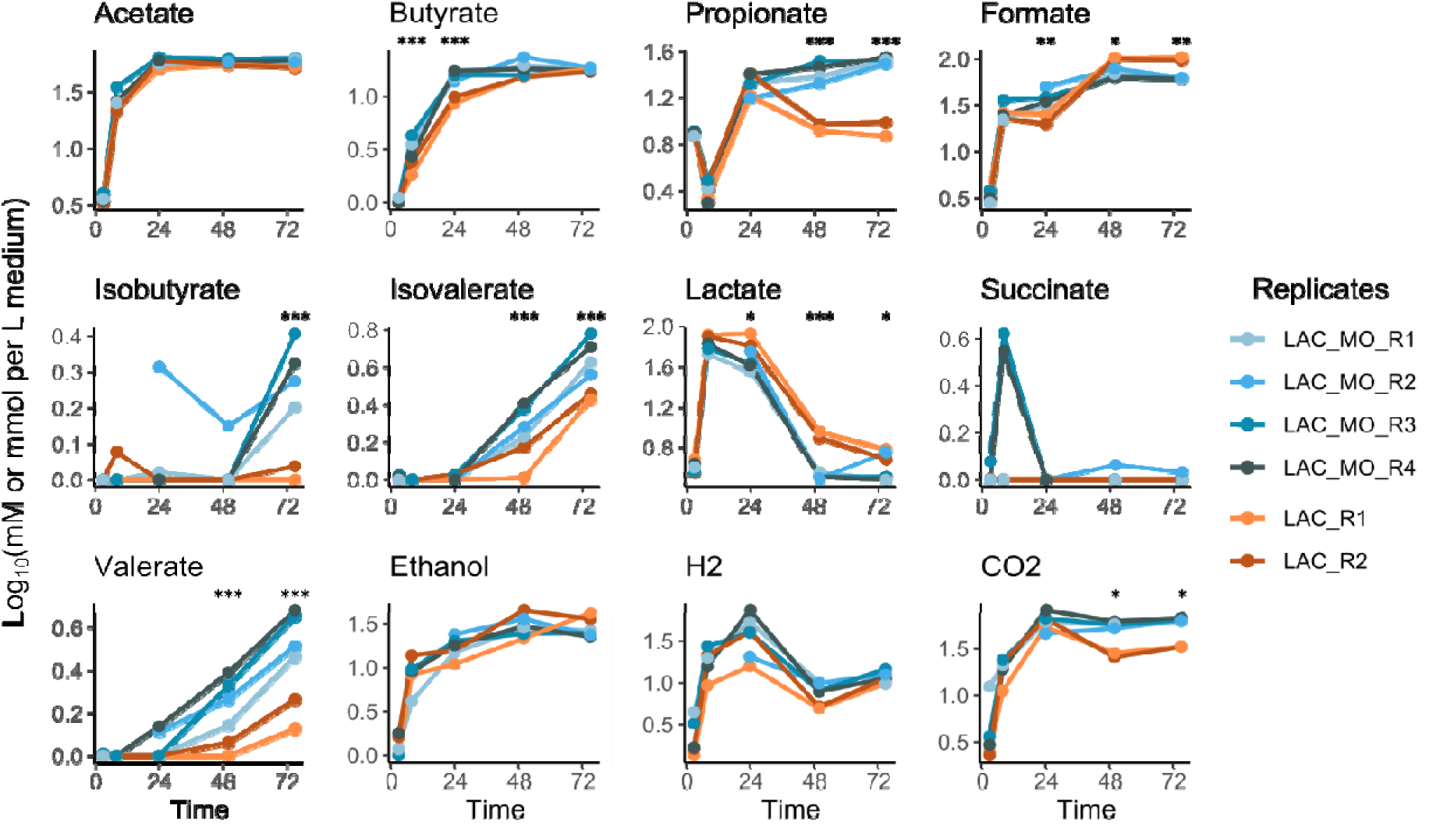
Metabolic transition from mixed-acid to SCFA fermentation enhanced by MO supplementation. Metabolite production in cultures of piglet fecal microbiota during batch phase (0-24 h) and chemostat phase (24-75 h) between replicas (R) of cultivations with lactose (LAC) and lactose-milk oligo-saccharide (LAC-MO) medium. Statistical significance is calculated between LAC-MO and LAC experiments.

### Fecal chemostat cultivation established a stable phage community devoid of eukaryotic viruses

When bacteria grow in a bioreactor inoculated with fecal matter, they engage in a dynamic interaction with the vast number of both virulent and temperate fecal phages^55^. To investigate the viral community fluctuations during fermentation, we isolated virus-like particles from the cultures (Fig. 1B) at each sample point followed by metagenomics sequencing. Relative to the fecal inoculum, the viral richnes decreased by 50% during the batch phase and further decreased during the chemostat phase, resulting in an overall decrease to 25% of the initial level (Fig 3A). The dilution occuring at inoculation (1:800 w/v) may explain the initial loss, as it might reduce low-abundant viruses to levels below the sequencing depth. In the chemostat phase, most phages with non- or slow-growing bacterial hosts were likely reduced alongside the eukaryotic viral elimination.

**Figure 3.**
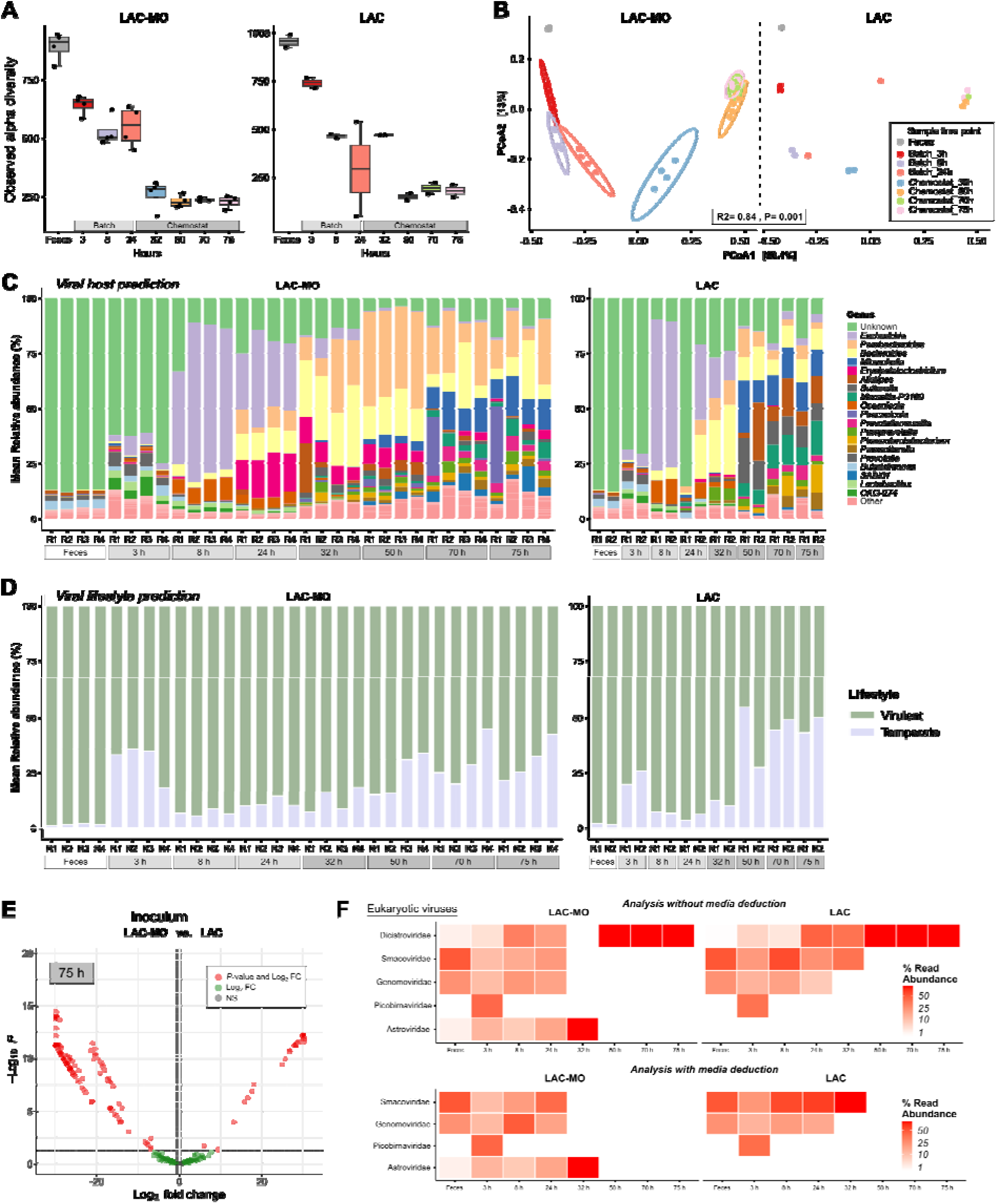
Chemostat cultutivation with lactose medium (LAC) or lactose-milk oligosaccharide medium (LAC-MO) yields selective but stable phage communities while depleting eukaryotic viruses. A) Number of observed viruses. B) Principal component analysis (PCoA) plot visualizing viral beta diversity over time, based on Bray-Curtis dissimilarity metrics. C) Relative abundance of viral bacterial hosts per replicate summarized at genus level. D) Predicted lifestyle of bacteriophages based on presence (temperate) or absence (virulent) of integrase and/or recombinase genes. E) Vulcano plot visualizing viral OTU differences between substrates at 75 hours. The DESeq2 enrichment analysis with FDR correction was based on viral metagenomics sequencing. Horizontal dotted lines show significance level of *P* < 0.05 and vertical lines show ± 0.6 log2 fold change (FC) cut-off. NS = non-significant. F) Depletion of eukaryotic viral families presented as heatmaps. The upper panel shows a raw analysis of culture samples. The lower panel shows the analysis, where the baseline viral content in the medium was deducted from culture samples. Color intensity indicates the relative abundance of specific eukaryotic viral contigs from total viral eukaryotic contigs.

This resulted in a final conserved community of around 200-250 viruses with similar levels for pure lactose cultures and MO-supplemented cultures. The community compositions changed dramatically during fermentation until reaching a stable level from five volume exchanges and onwards with high similarity between replicates (50-75 h, Fig. 3B), closely resembling the bacterial dynamics although without an obvious MO effect. When examining relative abundances, the majority of viruses were either classified as *Caudoviricetes* (class of tailed phages), *Microviridae* (family of small, non-tailed phages), or unknown viruses (Suppl. Fig. S3A). When predicting bacterial hosts, the fecal inoculum mainly harbored viruses infecting unknown bacteria (Fig. 3C). Phages targeting *Escherichia* began to increase after 8 hours, suggesting the induction of coliphages in response to the *Escherichia* expansion seen at 3 hours (Fig. 1D). In the chemostat phase, phages targeting *Bacteroides* and *Parabacteroides* started to emerge. These communities were best maintained during MO supplementation, where especially *Parabacteroides* phages reached relatively higher levels in end cultures. Interestingly, lifestyle prediction showed that the fecal inoculum mainly consisted of potential virulent phages, whereas the relative abundance of temperate phages steadily increased over time in the chemostat with no obvious effect of MO supplementation (Fig. 3D). Consequently, the transferred phages differed in terms of lifestyle between FVT and chemostat-propagated virome. DESeq2 analysis after 10 volume exchanges (Fig. 3E) revealed that adding MOs generally entailed a larger upregulation of specific viral OTUs compared to lactose only (134 vs. 27 vOTUs, respectively).

To uncover the success of eukaryotic viral elimination, eukaryotic vOTU members were manually identified and assessed in relative abundances over time (Fig. 3F). A total of five eukaryotic viral families were present in the fecal inoculum and batch phases. After five volume exchanges (50 h), four of these families were successfully eliminated, however, *Dicistroviridae* gradually increased during chemostat mode. Conflicting with our dilution hypothesis, we subsequently subtracted all viral contigs present in the medium and repeated the analysis. This caused *Dicistroviridae* to disappear, suggesting that this particular virus was continuously introduced via the medium rather than propagation. Hence, viral contigs present in the medium were deducted prior to all the *in vitro* viral analyses presented (Fig. 3). Conclusively, although less diverse than the native fecal virome, both chemostat settings created stable cultures of phages with an undetectable level of known eukaryotic viruses. The pronounced level of *Bacteroidota-*specific phages in MO-supplemented cultures revealed substrate-driven differences in phage host-specificities, worthy of further investigation.

### Fecal and chemostat-propagated viromes exhibit similar immunogenic potential

Virus-like particles (VLPs) were isolated from the feces (fecal virome transfer, FVT) and the 75-hour chemostats. CVT comprised of VLPs isolated from chemostats with lactose medium, and CVT-MO comprised of VLPs isolated from chemostats with lactose and MO medium (Fig. 4A). Any medium residuals including MOs were removed during this procedure (Fig. 1B). Chemostats with MOs produced end cultures with similar levels of VLPs compared to propagation with lactose (1.1-1.7×10^9^ and 1.1-1.2×10^9^ VLPs/mL, respectively). These liquid cultures had a more than 10-fold lower VLP level than raw feces (3.7×10^10^ VLPs/g). When preparing the intervention inocula, the dose was therefore adjusted to match similar absolute VLP levels (Fig. 4B) while carrying different levels of viral diversity (Fig. 4C). Aerobic and anaerobic cultivation showed no bacterial growth and nucleic acid extraction yielded extremely low DNA levels (0.0-0.4 ng/µl), with no amplification of 16S rRNA gene segments. Therefore, the inocula were considered bacteria-free.

**Figure 4.**
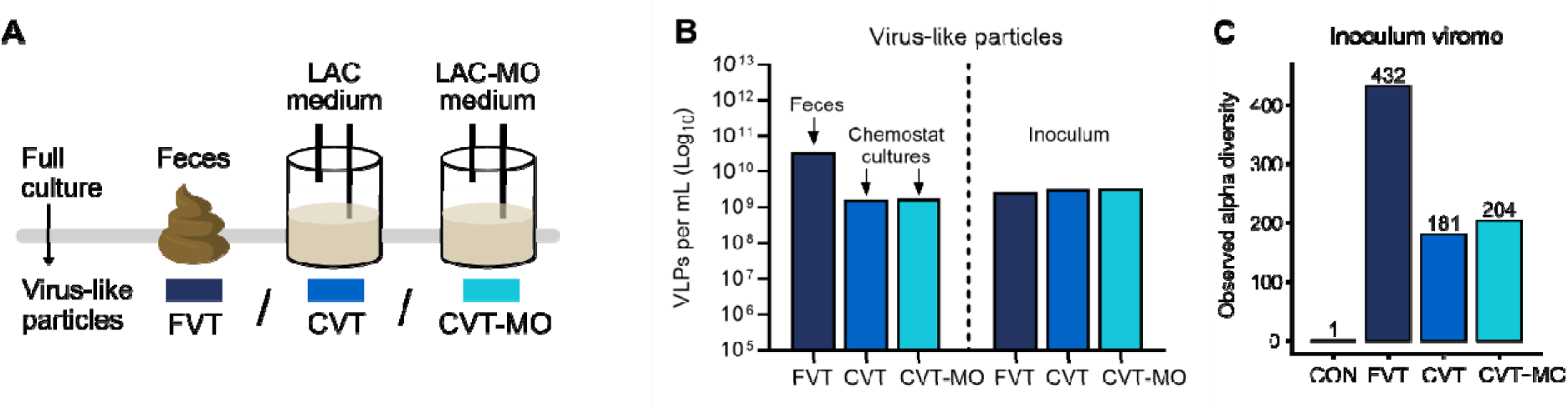
Preparation of bacteria-free virome inocula from feces and chemostats with matched virus levels. A) Inocula were produced by isolating virus-like particles (VLPs) from 1) donor feces to produce fecal virome transfer (FVT) solution, from 2) chemostat culture with lactose (LAC) medium to produce chemostat virome transfer (CVT) solution, and from 3) chemostat culture with lactose-milk oligosaccharide (LAC-MO) medium to produce CVT-MO solution. B) Estimation of average VLP level in donor feces (VLP/g), the chemostat cultures (VLP/mL), and the dose-adjusted inocula (VLP/mL). C) Number of observed viral species as a measure of viral alpha diversity in each inoculum and control (CON) solution.

A human monocyte cell line was employed to investigate the immunological impact of stimulation with the chemostat viromes. These responses were compared with the dose-adjusted native fecal virome (which includes eukaryotic viruses) and a dose-adjusted pure phage solution (free of eukaryotic viruses). Interestingly, chemostat viromes induced a lower expression of toll-like receptor (TLR) 7 compared to the native fecal virome (Suppl. Fig. S4). Since TLR-7 recognizes single-stranded RNA, this could reflect the reduced eukaryotic viral load in chemostat viromes. However, the significant expression mounted by the phage solution, consisting of double-stranded DNA phages, complicates the interpretation. Moreover, the typical virus-associated interferons were not markedly affected by the type of viral solution. Surprisingly, the chemostat viromes produced a higher expression of various pro- and anti-inflammatory cytokines, compared to the native fecal virome and with levels similar to the phage solution. This included tumor necrosis factor (TNF) α, IL-1β, IL-4, IL-8, IL-10, and transforming growth factor (TGF) β. Due to the endotoxin removal step and the use of a high MWCO during viral isolation, we consider it unlikely that these effects are driven by bacterial products in chemostat viromes. Given the obvious eukaryotic viral elimination, these data suggest that this innate immune cell line cannot fully distinguish between eukaryotic and prokaryotic viruses. Since fecal-derived eukaryotic viruses mainly infect intestinal cells, an *in vivo* stimulation may reveal a different response pattern.

### Chemostat-propagation mitigated side effects after virome transfer into preterm piglets

Previous studies revealed a good NEC protective effect of FVT but with simultaneous induction of early-onset diarrhea and weight loss in recipients^20^. Suspecting eukaryotic viruses to be the main driver, the chemostat method was applied. In *Piglet Experiment 1*, preterm piglets received a total dose of 10^10^ VLPs from feces or chemostat-propagated inoculums (FVT, CVT, CVT-MO) or a control solution shortly after birth (Fig. 5A). Three animals were euthanized early due to apnea or hindleg issues and were subsequently excluded from the experiment (1 CON, 1 FVT, and 1 CVT-MO). The remaining piglets (*n*=14-15/group) were followed for five days and displayed similar survival rates across groups (Fig. 5B). Upon euthanasia, gastrointestinal necropsy revealed an unusually low occurrence of NEC-like lesions in the control group as well as intervened animals (Fig. 5C-D). When investigating pathology incidences, the level of stomach necrosis was reduced in FVT pigs relative to controls (Suppl. Fig. S5A). Apart from this, incidences of hyperemia, hemorrhage, pneumatosis intestinalis, and necrosis were low and comparable between groups within each intestinal segment. Similarly, the histopathological assessment and tissue cytokine levels revealed no obvious differences (Fig. 5E and Suppl. Fig. S5B). Collectively, neither the FVT nor chemostat viromes appeared different from controls regarding NEC. However, the unfortunate lack of NEC-afflicted animals in the control group makes it impossible to draw firm conclusions in terms of treatment efficacy.

**Figure 5.**
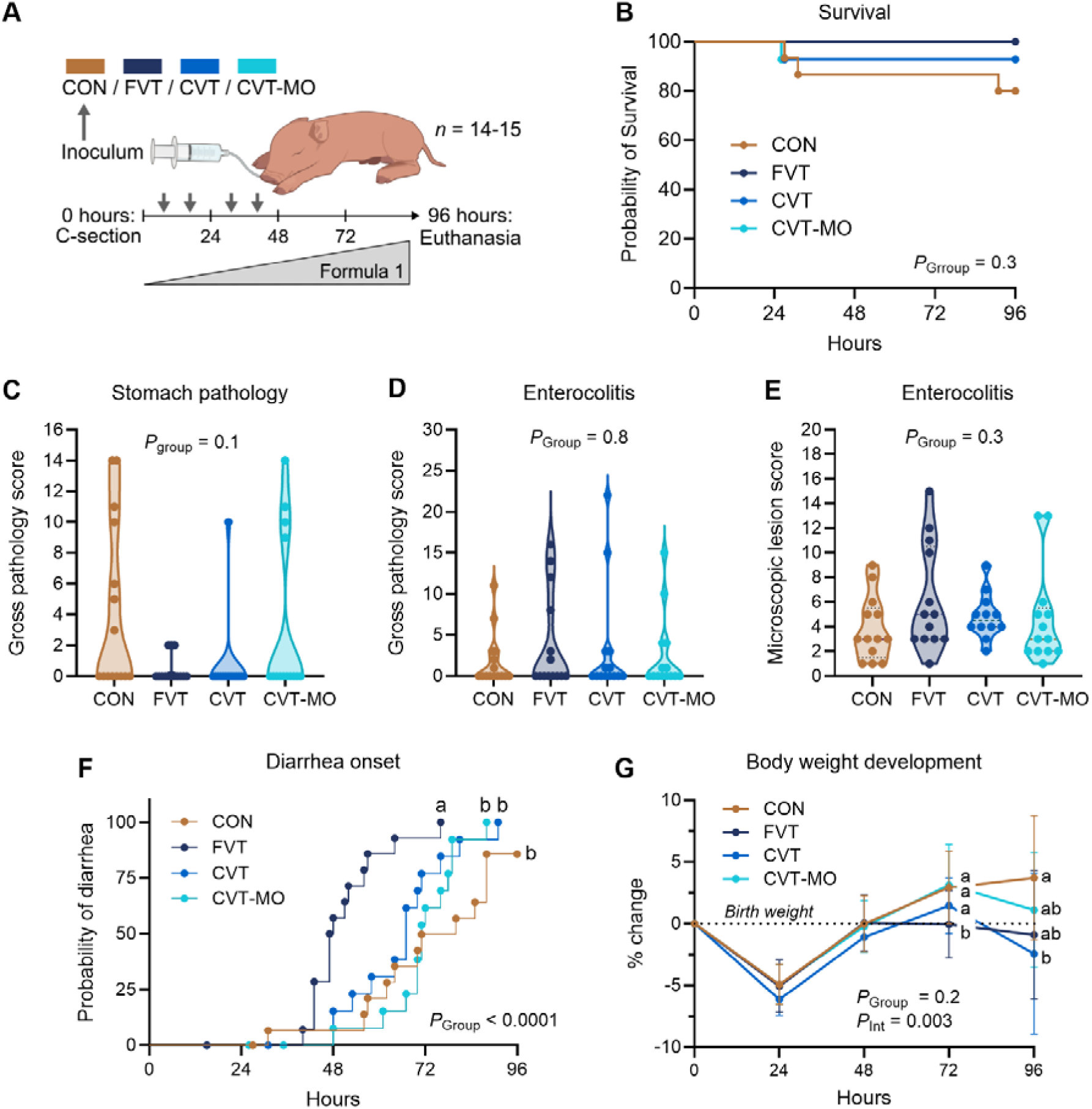
Chemostat propagation mitigates FVT-induced diarrhea but does not affect NEC-like lesioning in a low-incidence setting (*Piglet Experiment 1*). A) Cesarean section-delivered piglets received either control solution (CON) or one of the VLP-rich inocula (FVT, CVT, CVT-MO) over four oral treatments. The pigs were fed increasing volumes of enteral formula 1 until euthanasia. B) Survival curve. C) Gross pathology score in stomachs. D) Cumulative gross pathology score in small intestines and colons. E) Cumulative microscopic lesion score in small intestines and colons. F) Time to diarrhea onset. G) Body weight development showing percentage change from the median birth weight (±SD). Lines not sharing the same letter are significantly different (*P* < 0.05). *P*_Group_ = *P*-value for group effect*, P*_Int_ = *P*-value for group and time interaction. FVT = fecal virome transfer, CVT = chemostat virome transfer, and CVT-MO = CVT propagated with milk oligosaccharides (*n* = 14-15/group).

The clinical monitoring, however, revealed clear differences in the side effect pattern throughout the study. Replicating previous results^20^, FVT animals debuted with earlier diarrhea, leading to a reduced weight gain on day four (Fig. 5F-G). Both chemostat viromes were statistically similar to controls regarding diarrhea onset, suggesting the method successfully alleviated the effect of the diarrhea-inducing agent (Fig. 5F). By visual inspection, the diarrhea incidence of piglets receiving lactose or MO-propagated viromes appeared slightly accelerated, resulting in a decline in body weight by study termination in the CVT group (Fig. 5G). By day five, circulating neutrophil levels were trending higher in FVT animals compared to controls (*P* = 0.08, Suppl. Fig. S6A). On the contrary, FVT lymphocyte levels remained lower from day three and onwards. Although speculative, this implies that FVT may induce migration of lymphocytes to the intestines as well as granulopoiesis, resulting in a subsequent increase in circulating neutrophils. This response was mitigated by chemostat propagation. Consistent with previous findings, the FVT side effects were linked to small intestinal atrophy and reduced lactase activity in the proximal small intestine (Suppl. Fig. S6B-C). Here, the lactose-propagated virome entailed atrophy while the MO-propagated virome reduced lactase activity. Additionally, the lactulose-to-mannitol ratio increased only significantly in FVT animals, suggesting increased intestinal permeability by this treatment alone (Suppl. Fig. S6D). Conclusively, the side effects associated with fecal virome transfer were significantly improved, but not fully abolished, using the *in vitro* propagated chemostat viromes.

Gut virus engraftment following fecal virome transfer is lost after chemostat propagation Having demonstrated a difference in the clinical side effect pattern in response to exogenous native and chemostat-propagated fecal viruses, we next investigated the changes to the newborn recipient gut virome using viral metagenomics sequencing. We observed a modest increase in viral richness in FVT recipients relative to those receiving lactose-propagated fecal virome (Fig. 6A). However, changes to viral richness did not appear related with the occurrence of NEC (Fig. 6B). Interestingly, Bray-Curtis ordination plot clearly showed an alteration of the gut viral composition of the FVT group relative to all remaining groups, which were not different from one another (Fig. 6C). The binned taxonomic annotation showed mainly *Caudoviricetes* and a significant proportion of unannotated viruses across all groups (fig. 6D). In pairwise comparisons of single viral OTU relative abundances between treatment groups and control by DESeq2 analysis, we found a pattern of modest virus enrichment (32 enriched vs. 16 depleted) in the FVT group but neither of the chemostat virome groups (CVT: 9 vs. 16; CVT-MO: 17 vs. 15). All of these were annotated as unknown viruses or the class *Caudoviricetes*. Based on this viral enrichment analysis, we found no clear evidence of eukaryotic virus shedding to explain the clinical side effects of FVT as virus-induced. Importantly, when binning the predicted bacterial hosts, gut phages of the FVT recipients had a much greater specificity for *Lactobacillaceae* accounting for up to 50% relative abundance, whereas the remaining groups had hardly any *Lactobacillaceae* phages (Fig. 6E). Moreover, when looking into the engraftment potential of the transferred phages, assessed as the percentage of viral OTUs shared between group inoculum and recipient gut sample, only the FVT group showed an increase, whereas recipients of both chemostat-propagated viromes had levels similar to controls (Fig. 6F). Taken together, we documented that transfer of fecal virome leads to an engraftment of viruses short-term and that most of these are phages with specificity for bacteria in the *Lactobacillaceae* family. Chemostat propagation of fecal viruses prior to transplantation disrupted this engraftment.

**Figure 6.**
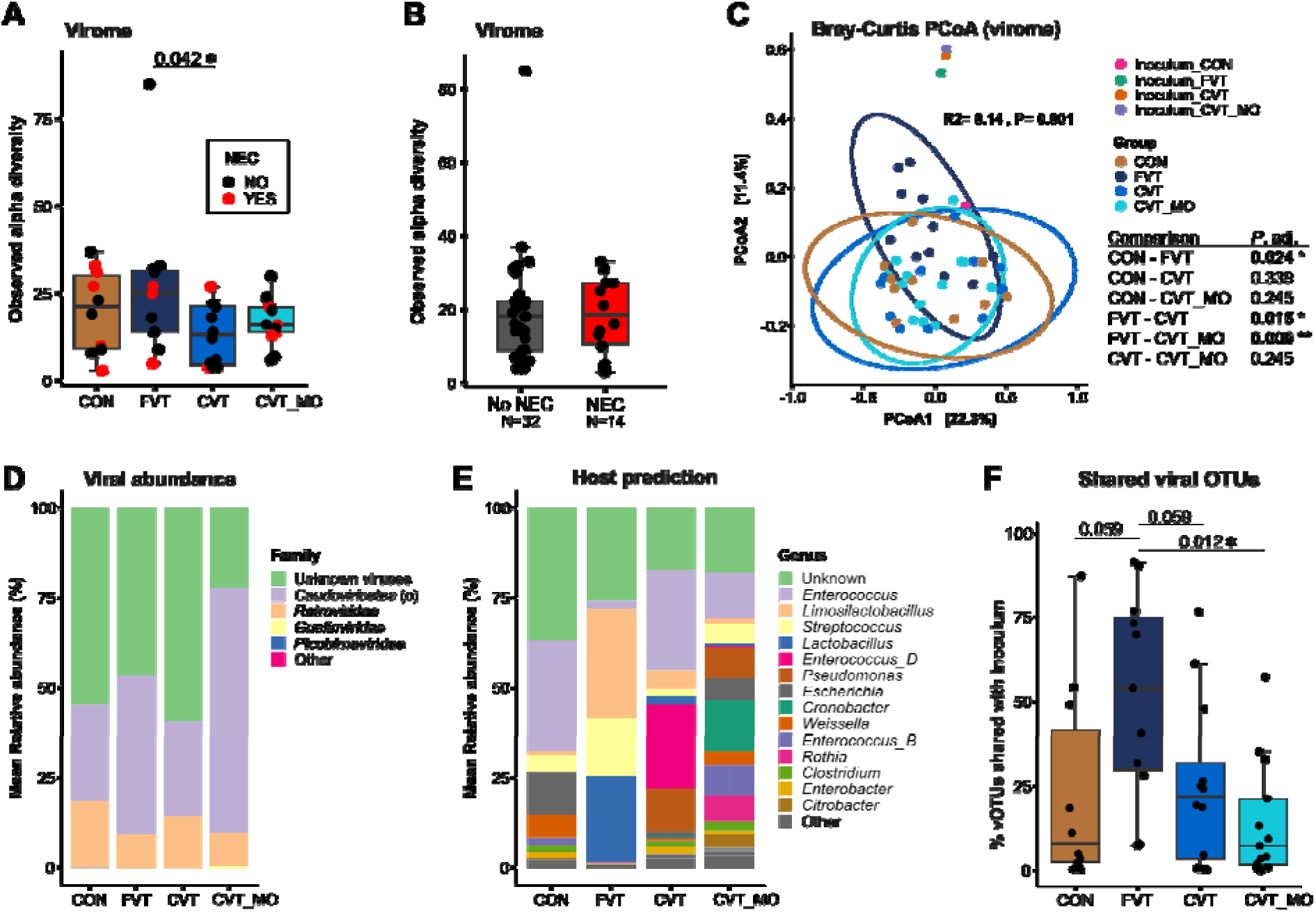
Native but not chemostat-propagated viromes reshape gut virome and enrich *Lactobacillaceae* phages (*Piglet Experiment* 1).. A-B) Number of observed viral species as a measure of viral alpha diversity between groups (A) and between NEC and healthy appearing colons (B). C) Principal coordinate analysis (PCoA) plot visualizing viral beta diversity based on Bray-Curtis dissimilarity metrics. False discovery rate (FDR) adjusted *P*-values for pairwise comparisons are reported in the adjacent table. D) Mean relative abundance of viral bacterial hosts summarized at genus level. E) Predicted lifestyle of bacteriophages based on presence (temperate) or absence (virulent) of integrase and/or recombinase genes. F) The percentage of shared viral OTUs between recipients of each group and the inoculum material of the group. CON = control, FVT = fecal virome transfer, CVT = chemostat virome transfer, and CVT-MO = CVT propagated with milk oligosaccharides (*n* = 10-13/group). \**P* < 0.05, \*\**P* < 0.01

Lactobacillaceae expansion following fecal virome transfer is lost after chemostat propagation The few animals with NEC (here defined as the presence of hemorrhage, pneumatosis intestinalis, and/or necrosis) had a higher bacterial richness in the colon, whereas no differences were seen between treatment groups (Fig. 7A-B). The bacterial composition in recipients of native FVT and MO-propagated fecal virome were slightly different from controls (Fig. 7C). From the taxonomic overview showing the most abundant taxa across groups, the most obvious difference between virome treatment groups and control was an increase in the relative abundance of *Streptococcus,* the same bacterial genus that dominated the chemostat community (Fig. 7D). Additionally, only FVT recipients harboured an increased proportion of *Lactobacillus amylovorus* (Fig. 7D). We performed pairwise DESeq2 analysis to identify all enriched and depleted bacterial species between treatment groups and control. Notably, FVT recipients were the only ones enriched with three different species of *Lactobacillaceae*, and concomitantly depleted in the archetypical NEC pathogen *Clostridium perfringens* (Suppl. Fig. S7 and Suppl. Table S3)^56^. The compositional effects of MO-propagated fecal virome were also evident by DESeq2, where a simultaneous reduction in both *Lactobacillaceae* and *Clostridium perfringens* was observed.

**Figure 7.**
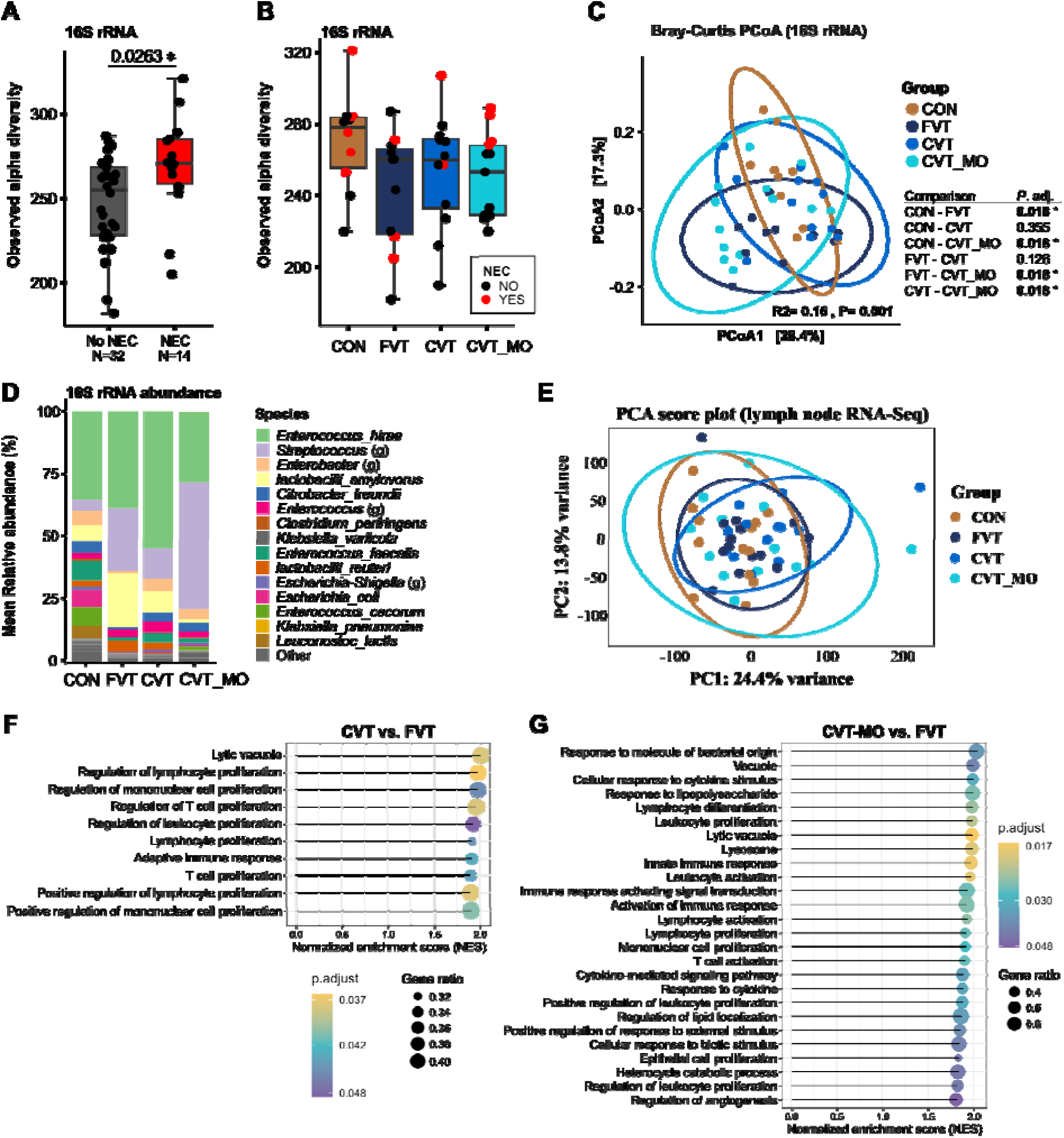
Virome treatments induce modest changes in the gut bacteriome, but have a limited impact on mesenteric lymph node gene expression (*Piglet Experiment 1*). A-B) Number of observed bacterial species between NEC and healthy appearing colons (A) and between groups (B). C) Principal coordinate analysis (PCoA) plot visualizing bacterial compositional differences as determined by Bray-Curtis dissimilarity metrics. False discovery rate (FDR) adjusted *P*-values for pairwise comparisons are reported in the adjacent table. D) Mean relative bacterial abundance summarized at species or genus (g) level. E) Principal component analysis (PCA) score plot illustrating the global gene expression differences in mesenteric lymph nodes. F-G) Gene set enrichment analysis (GSEA) based on gene ontology classification, showing all significantly enriched pathways (false discovery rate adjusted P-value < 0.05) between CVT and FVT (F) and between CVT-MO and FVT (G). The size of the dots indicates the gene ratio, while the yellow color indicates a lower adjusted P-value. NS = non-significant. CON = control, FVT = fecal virome transfer, CVT = chemostat virome transfer, and CVT-MO = CVT propagated with milk oligosaccharides (*n* = 10-13/group). \**P* < 0.05.

To investigate a potential link between gut microbes and mammalian host response further, we performed bulk RNA-Seq on mesenteric lymph nodes. We observed no major shifts in global gene expression patterns based on a principal component analysis (Fig. 7E). Gene set enrichment analyses did however identify a list of Gene Ontology pathways almost exclusively involved in immune cell activation. Contrary to expectations, these were all enriched in both groups receiving chemostat-propagated virome relative to FVT (Fig. 7F-G). As such, this finding does not support a eukaryotic viral infection by the transferred native fecal virome as an explanation for the observed side effects. However, the collected lymph nodes were draining more distal intestinal segments (near the ileocolic artery), far from the site where donor viruses were first introduced by oral gavage. Since FVT affects the small intestine (Suppl. Fig. S6B-D), sampling from more proximal regions (near the superior mesenteric artery) may have revealed different treatment responses. Additionally, sampling at the time of FVT-associated diarrhea acceleration (48 hours) might uncover group differences not apparent at euthanasia (96 hours), when all groups displayed similar diarrhea levels.

### Milk oligosaccharide-propagated fecal virome did not prevent NEC-like lesioning

The unusually low levels of gut lesions among controls in *Piglet Experiment 1* did not allow for the investigation of a potential NEC-reducing effect of chemostat-propagated fecal viruses. This prompted *Piglet Experiment 2*, where two litters of pigs were fed a UHT-treated infant formula (formula 2) with a demonstrated higher NEC-inducing capacity (Fig. 8A)^37^. To minimize animal numbers, we focused on investigating the MO-propagated virome, as this treatment had the greatest microbiome impact of the chemostat-propagated viromes. Both groups displayed similar survival rates throughout the study period (Fig. 8B). At necropsy, the general level of intestinal gross pathology was markedly higher compared to *Piglet Experiment 1* (median score of 7 vs. 0). Chemostat virome recipients exhibited more severe stomach lesions (Fig. 8C and Suppl. Fig. S8A), not previously noted. The overall enterocolitis level was similar at macroscopic and microscopic assessments (Fig. 8D-E). However, the chemostat virome group showed a slightly higher incidence of small intestinal bleeding (*P* = 0.05, Suppl. Fig. S8B) but with subtly lower IL-8 levels (*P* = 0.03, Suppl. Fig. S8D). The visual trends of slightly earlier diarrhea and decreased body weight aligned with those observed in *Piglet Experiment 1* (Fig. 8F-G). Again, these trends were not statistically significant and did not entail intestinal atrophy (Suppl. Fig. S8E). However, intestinal permeability was slightly increased (*P* = 0.01, Suppl. Fig. S8F). The colonic bacterial analysis showed only minor changes, consistent with the previous findings (Suppl. Fig. S9). Collectively, the MO-propagated fecal virome caused gastritis and appeared to mildly compromise small intestinal health. This highlights that while chemostat propagation effectively mitigates virus-associated diarrhea, the method needs further optimization to target NEC.

**Figure 8.**
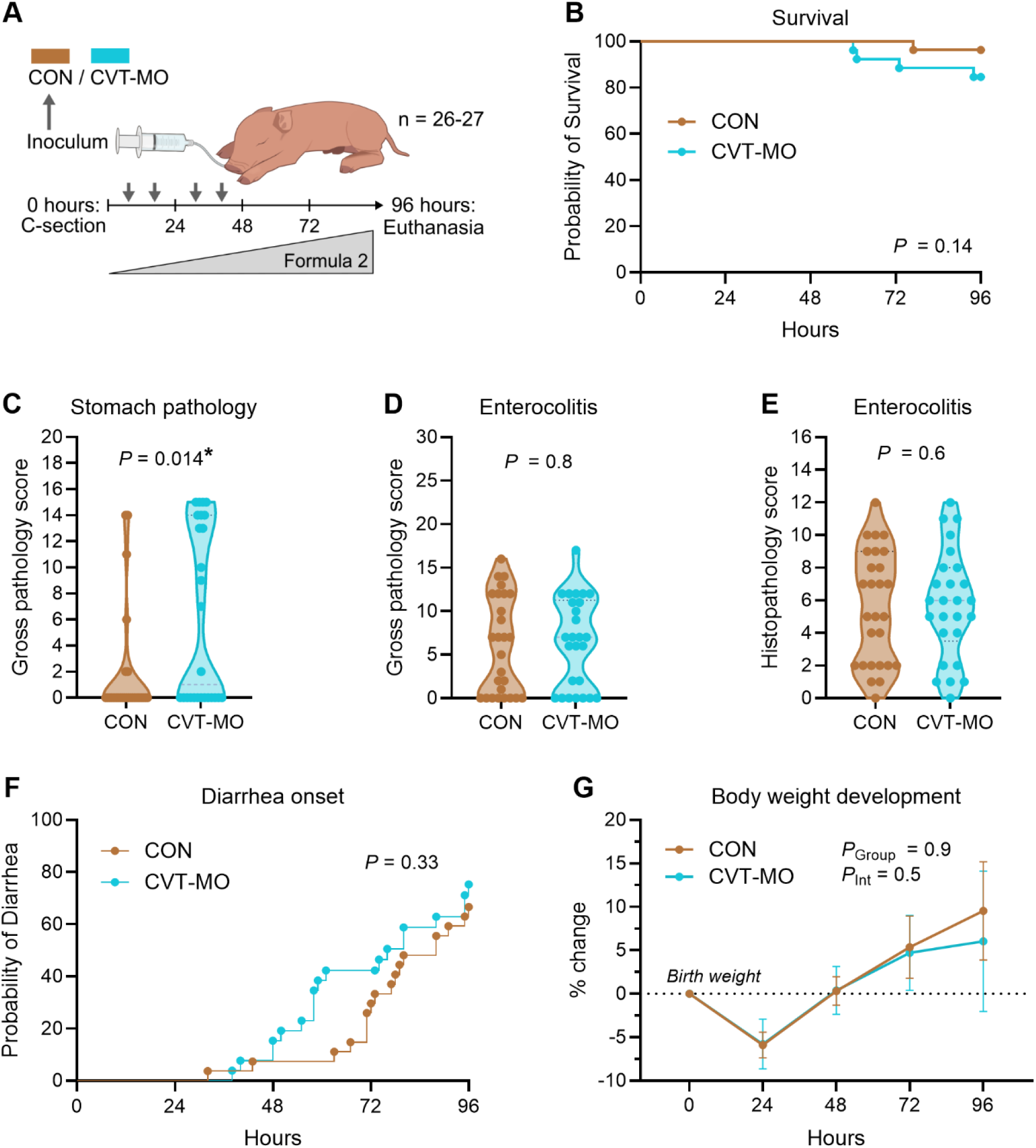
MO-propagated virome modestly exacerbates gastric injury and fails to improve NEC (*Piglet Experiment 2*). A) Overview of study design. Cesarean section-delivered piglets were allocated to one control group (CON) receiving SM buffer and one treatment group receiving chemostat virome transfer propagated with milk oligosaccharides (CVT-MO). The pigs were fed increasing volumes of enteral formula 2 until euthanasia after 96 hours. B) Survival curve. C) Gross pathology score in stomachs. D) Cumulative gross pathology score in small intestines and colons. E) Cumulative microscopic lesion score in small intestines and colons. F) Time to diarrhea onset. G) Body weight development showing percentage change from the median birth weight (±SD). *P*_Group_ = *P*-value for group effect*, P*_Int_ = *P*-value for group and time interaction. \**P* < 0.05 (*n* = 26-27/group).

## Discussion

Transfer of fecal phages has shown promise in modulating the preterm gut microbiome and reducing NEC under experimental settings^17,20^. However, as diarrhea occurs simultaneously^20^, this highlights a need for further refinements to increase the level of safety while maintaining efficacy. Expanding on previous research^24^, we investigated whether the elimination of eukaryotic viruses during chemostat processing could reduce potential side effects of fecal virome transfers for preterm neonates while preserving a phage composition capable of correcting GM dysbiosis and preventing NEC.

### Choice of parameters for chemostat propagation

We have previously demonstrated the anaerobic chemostat setup as a method to cultivate stable fecal bacterial and viral communities from adult human and murine donors^22,24^. To adapt this for cultivating colon and cecal content from 10-day-old suckling piglets, several adjustments were required. First and foremost, the growth medium substrates were adjusted to closely resemble a chemostat setup mimicking the proximal colon of breastfed neonates^23^, which contained lactose as the main carbohydrate source along with gastric mucin. However, since most lactose is absorbed before reaching the colon^57^, we considered it important to investigate the effect of MOs as an additional substrate for cultivation of a milk-shaped fecal microbiome^58^. Despite being less diverse than human milk, porcine milk contains 30-50 different MOs, several of which are also found in human milk^32,53,54^. For simplicity, we selected three key MOs found in both species (2’FL, 3’SL, and LNnT) and known to be degradable by piglet fecal bacteria^59^. Opposite to infants, young pigs harbor very few bifidobacteria^60^. Therefore, the gradual emergence of fecal *Bacteroidota* species during chemostat effectively demonstrated the MO-utilizing potential of these gut bacteria inhabiting healthy suckling piglets^61,62^. MOs allowed the cultivation of around 150 additional bacterial species and increased the bacterial similarity to better resemble the donor. Optimizing the MO composition may promote the dominance of more favorable MO-degrading *Bacteroides* spp. over *B. pyogenes*^63^. Together, this provides a strong proof-of-concept for using MOs to culture neonatal fecal bacteria *in vitro*.

Despite the restorative effect of MO supplementation, the bacterial composition of the end culture was fundamentally different from the donor feces, which was primarily due to losing virtually all lactobacilli. As lactobacilli are fast-growing bacteria that primarily utilize lactose^64^, which was present in ample amounts in both chemostats (especially LAC), the growth inferiority is unlikely due to substrate limitations. Instead, it may be due to suboptimal pH levels. Studies report varying colonic pH levels in suckling pigs ranging from 6.3 to 7.2^65–67^ and thus an intermediate level of 6.5 was chosen for the experiment. The study by Pham *et al*. investigated pH levels ranging from 5 to 7 and found infant lactobacilli showed optimal growth at pH 5, whereas *Bacteroides* preferred pH 6-7^23^. Alongside the current findings, this suggests that lower pH levels are needed to support the growth of both *Lactobacillaceae* and *Bacteroides* (and bifidobacteria) in future neonatal chemostats.

Chemostat culturing of neonatal feces using the presented parameters resulted in a significant loss in viral richness, which was not remedied by the addition of MO. Our previous study using adult fecal material indicated higher phage preservation at slower dilution rates (0.05 h^-1^) for murine, but not for human fecal cultures^24^. As the pig GIT resembles humans more than mice^68^, we thus decided on the higher dilution rate (0.2 h^-1^) to ensure a significant eukaryotic viral elimination. Albeit, the current study suggests that slower dilution may be required for proper *in vitro* phage propagation of a neonatal gut virome. The viral inoculation dose of 10^10^ VLPs/kg was chosen to match previous studies demonstrating NEC prevention (ref to ISME + gut microbes paper). However, if a transferred phage encounters a suitable host and enters the lytic cycle, phage titers may increase drastically. Consequently, greater phage richness will, irrespective of the starting dose, increase the probability of productive phage-bacterial interactions and their overall impact on the gut microbiome.

### Correction of preterm gut dysbiosis

Although the chemostats poorly resembled the piglet gut microbiome, the MO-induced increase in *Bacteroidota*-specific phages could beneficially modulate the recipient gut microbiome. Indeed, we found that a significant proportion of phages in the chemostat virome, including those targeting *Bacteroidota,* were induced prophages, as opposed to the fecal inoculum containing almost exclusively virulent phages. Although less abundant in human infants compared to piglets, *Bacteroides* is a part of the normal developing microbiome^11,69,70^ and is positively linked to intestinal health^71^. FVT has repeatedly been shown to increase the similarity between donor and recipient bacterial compositions, without administering the actual donor bacteria^72,73^. We have previously observed increased levels of *Bacteroides* spp. in piglet colons after receiving NEC-reducing FMT or FVT treatments^17,74^. However, the MO-propagated chemostat virome did not increase *Bacteroides* or *Parabacteroides* spp. in the recipient’s gut, nor did it provide NEC protection when tested in a high-incidence setting in *Experiment 2*. We find it likely that this lack of engraftment could be due to an absence of bacterial hosts in the preterm piglet recipients.

As Experiment 1 piglets had unusually low levels of intestinal lesions, the NEC-reducing capacity of this particular donor fecal virome remains unknown, although the FVT concept has been proven efficacious on previous occasions using similar donor material^17,20^. Importantly, the virome transfer was capable of reducing Clostridium perfringens while increasing the abundance of Lactobacillaceae members. Although NEC cases were sparse, these showed the opposite trend regarding these bacteria, thus indicating a restorative effect of FVT on NEC-associated dysbiosis. Peculiarly, viral metagenomics data indicated that mainly virulent phages from the FVT inoculum engrafted in recipients and that these had Lactobacillaceae as hosts. This contrasts a simple lytic predation of target bacteria as an explanatory mechanism. This virulent classification, however, only reflects the absence of known recombinase or integrase genes, meaning these phages are considered potentially virulent based on current annotations. Regardless, the lack of Lactobacillaceae in chemostats could potentially explain the lack of recipient gut engraftment of the chemostat-propagated viromes despite transferring similar numbers of virus-like particles. As we have previously seen Lactobacillaceae abundance linked to NEC protection^30,74^, this again highlights a target for optimizing chemostat conditions to maintain this bacterial family, thereby enabling efficacy of chemostat propagated feces. Moreover, whether the reduction of C. perfringens was due to transferred lytic phages or as a result of competitive exclusion by other bacteria remains unknown. However, we acknowledge that the absence of demonstrated NEC prevention by the native donor virome is a significant limitation to conclusions regarding efficacy. Without a protective baseline, we cannot assess whether the virome efficacy was lost during chemostat propagation. Consequently, we cannot confirm or dismiss the hypothesis that chemostats can preserve a phage community capable of preventing NEC.

### Transfer of chemostat viromes entails later diarrhea

We have previously demonstrated the use of chemostat viromes to improve the health status of mice during metabolic syndrome and *Clostridioides difficile* infection^26,27^. This initiative aimed to 1) overcome variations in efficacy between FVT donors and 2) enhance safety by removing donor-derived eukaryotic viruses^24^. Since the existing studies showed no side effects, the importance of eukaryotic virus removal for improved FVT safety has remained a theoretical concern. The present study, however, showed that FVT induces diarrhea consistently and that chemostat propagation delays diarrhea onset. Regardless, since the viral metagenomics analysis did not identify a specific eukaryotic viral agent, the exact mechanism for how chemostat propagation reduces side effects remains elusive^20^. The four eukaryotic viral families found in feces or batch cultures (*Smacoviridae, Genomoviridae, Picobirnaviridae*, and *Astroviridae*) were not found upregulated in FVT recipients, nor was any inflammatory signal detected in mesenteric lymph nodes on day five. As diarrhea appeared on days 2-3 alongside reduced systemic lymphocyte counts, earlier analyses may have revealed different trends between groups.

## Conclusion

In conclusion, we show that chemostat propagated fecal virome reduces eukaryotic virus abundance drastically and minimizes gastrointestinal side effects when transferred into newborn recipients. Chemostat cultivation is a promising method to generate complex but reproducible fecal viromes, and supplementing the culture with MOs shows promise for approximating milk-shaped fecal microbiomes. However, as neither the chemostat-propagated viromes nor the native donor virome demonstrated NEC prevention, the efficacy of the chemostat approach remains inconclusive. MO composition, pH, and dilution rate may need optimization for better virus engraftment, recipient microbiome restoration, and NEC protection in future studies.

## Author’s contributions

SMO, SA, TR, DSN, AB, and KA conceived and planned the experiments. SMO, SA, and KA performed the chemostat experiments. SA and KA performed metabolite analyses. SMO, MS, and AB performed the animal experiments. XM carried out bacteriome analyses. XM and FL carried out virome analyses. JZ performed additional lymph node transcriptome analyses. All authors were involved in critical evaluation and interpretation of results. SMO and AB drafted the manuscript with inputs from all the other authors.

## Availability of data and materials

All data associated with this study are present in the paper or the Supplementary Materials. All sequencing datasets are available on the BioProject database under accession number PRJNA1121201.

