## Supplementary material for "Chemostat culturing reduces fecal eukaryotic virus load and delays diarrhea after virome transplantation"

**Supplementary Figures**

Lactose medium (LAC)

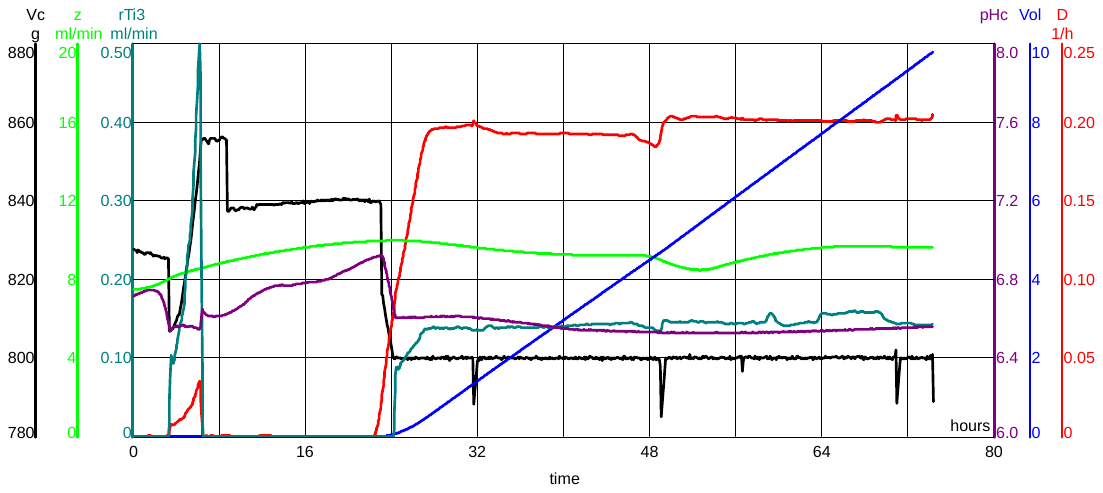

Lactose and milk oligosaccharide medium (LAC-MO)
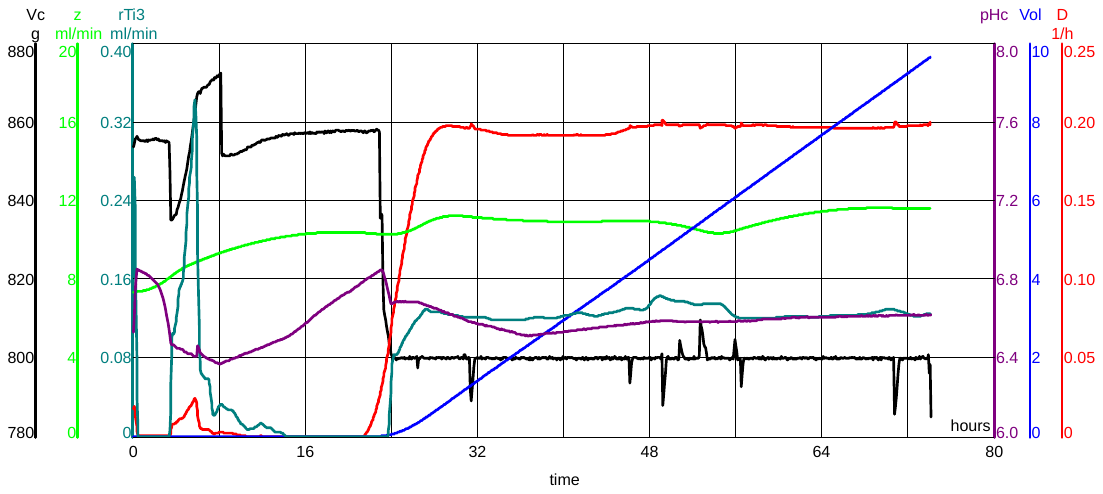

**Supplementary** Figure S1. Dynamics of experimental parameters during chemostat cultivation (D = 0.2 1/h) of pig fecal cultures. Replicate 2 from each medium condition is shown. rTi2 – titration rate of 3 M NaOH (mL/min), Vc – fermenter volume, z – microbial gas production rate (ml/min) calculated as the difference of GasS-Gas, where GasS – total gas flow rate (N2+microbial), Gas – amount (mL) of nitrogen gas flown through the fermenter, D – dilution rate (1/h), Vol – indicates volume exchanges.

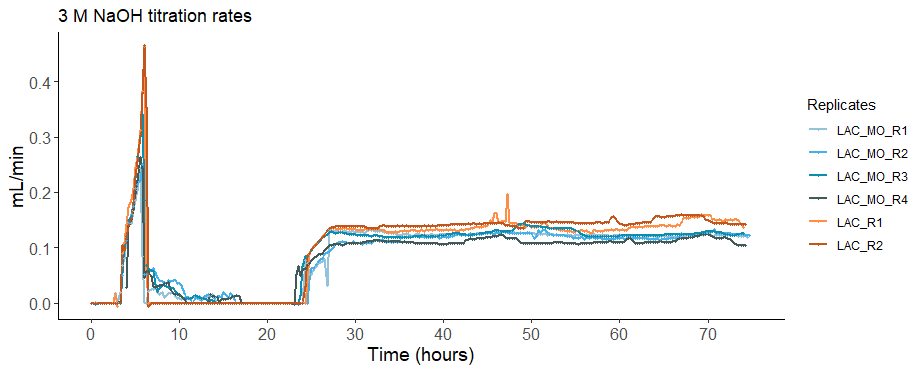

**Supplementary** Figure S2. The titration rate of 3 M NaOH to maintain pH during the batch phase (0-24 h) and chemostat phase (24-75 h) between replicas (R) of chemostats with lactose (LAC) and lactose-milk oligosaccharide (LAC-MO) medium.

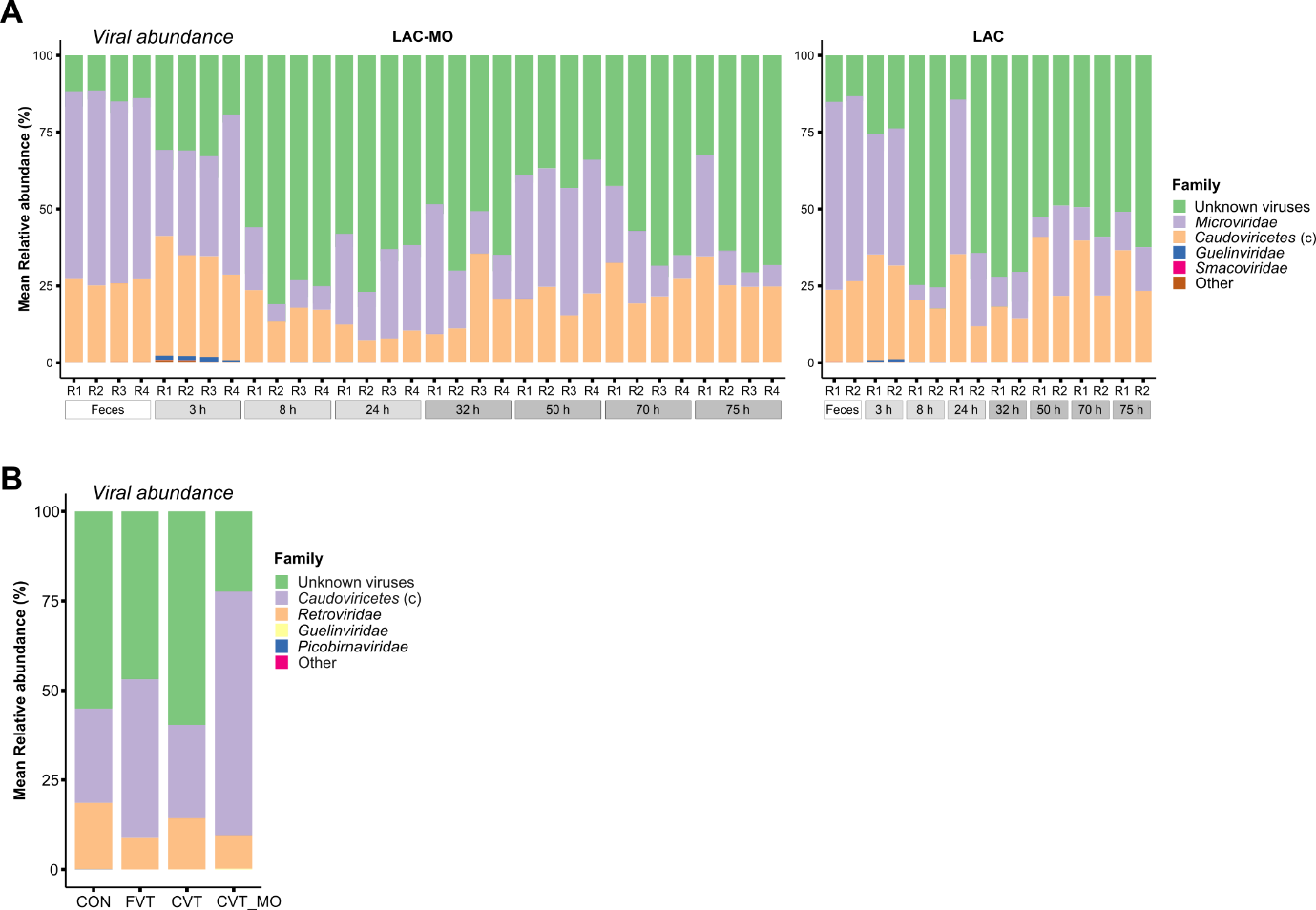

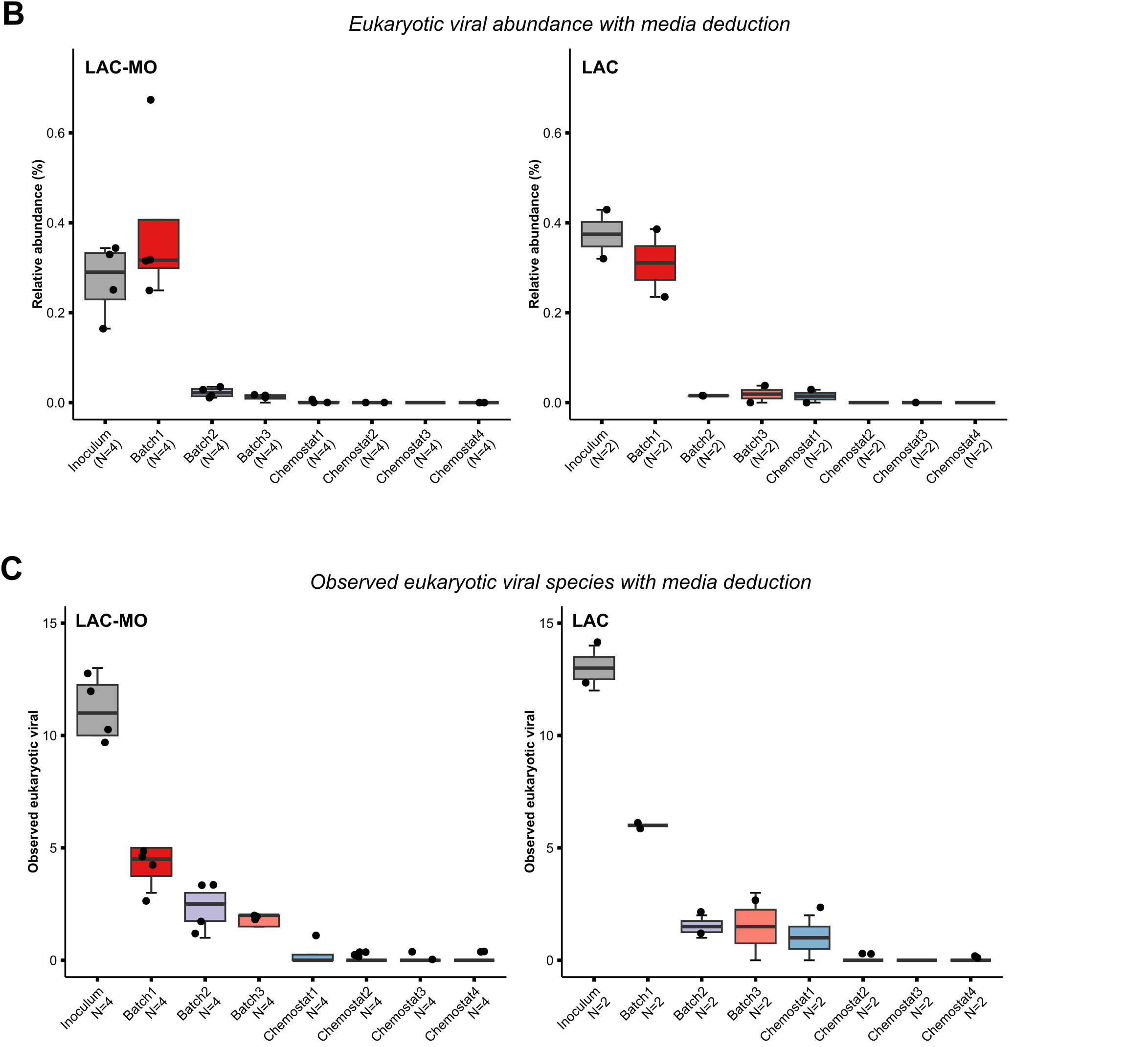

**Supplementary figure S3**. Viral development in the chemostat process. A) Overall virome compositional development in chemostat replicates with lactose medium (LAC) or lactose-milk oligosaccharide medium (LAC-MO). Shows relative viral abundance per replicate (R) summarized at family or class (c) level. B) Eukaryotic viral development in the same chemostat replicates after deduction of the media virome. C) Numbers of observed eukaryotic viral species in chemostat replicates after deduction of the media virome.

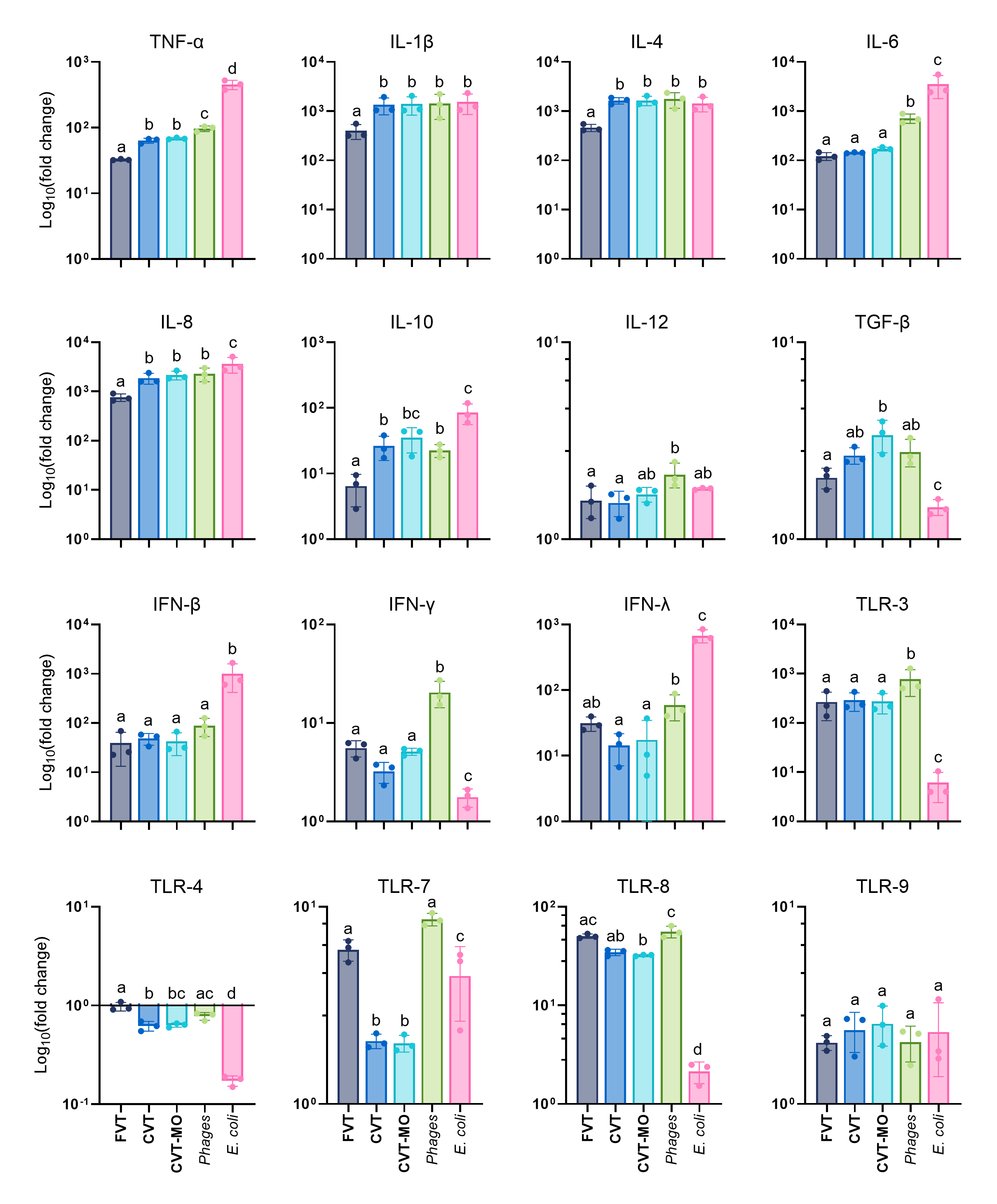

**Supplementary** Figure S4. Gene expression of THP-1 cells stimulated with a pure phage solution (*Phages*), *E. coli* bacteria, fecal virome solution (FVT), or chemostat virome solutions; CVT comprised virus-like particles (VLPs) isolated from chemostats with LAC medium and CVT-MO comprised VLPs isolated from chemostats with LAC-MO medium. Expression was calculated as fold changes relative to the level of expression in cells stimulated with autoclaved SM buffer. Bars not sharing the same letter differ significantly (*P* < 0.05). TNF = tumor necrosis factor, IL = interleukin, TFG = transforming growth factor, IFN = interferon, TLR = toll-like receptor.

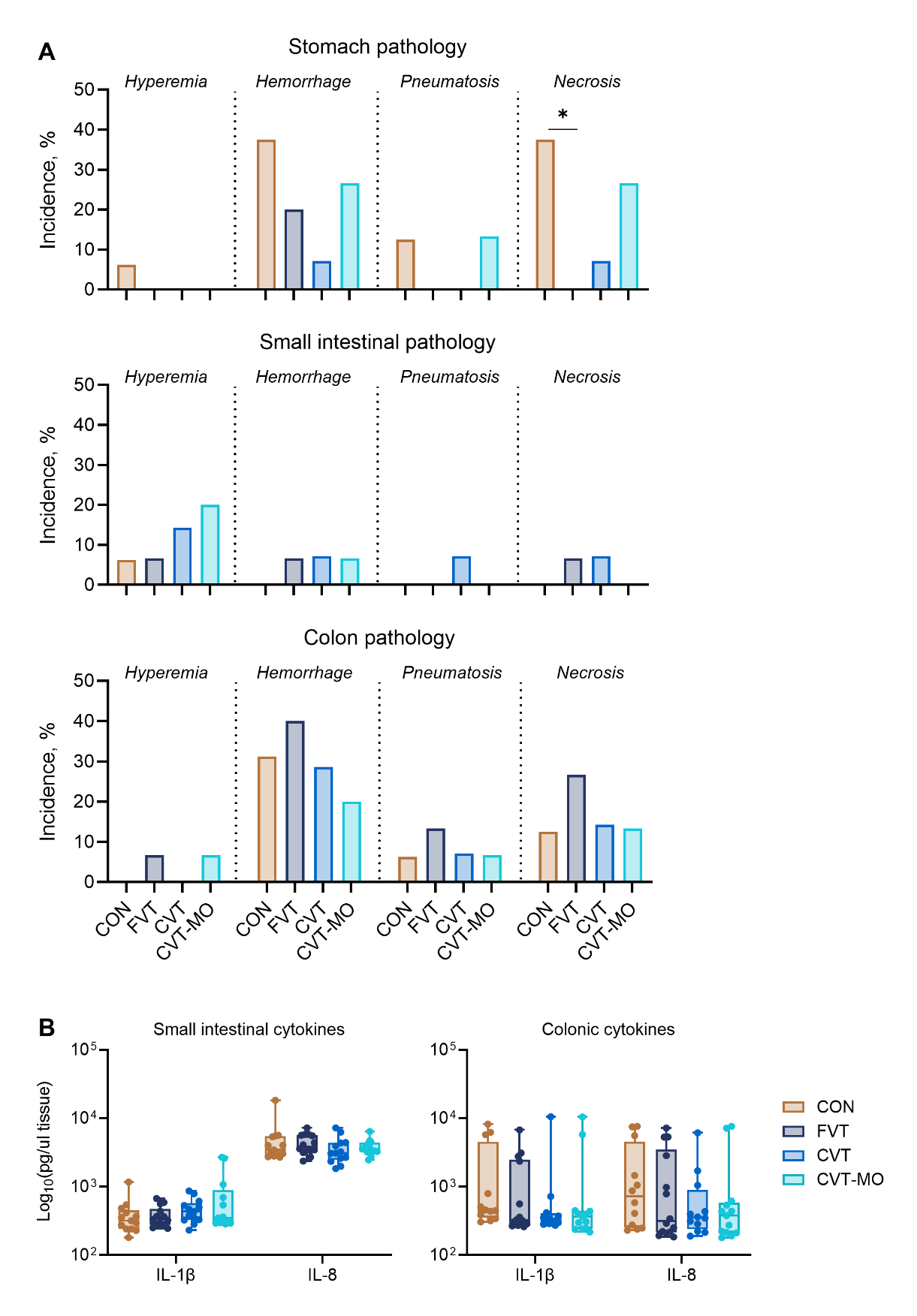

**Supplementary** Figure S5**.** Supporting data for assessment of intestinal pathology. A) Incidences of hyperemia, hemorrhage, pneumatosis intestinalis (intramural gas), and necrosis. B) Interleukin (IL) 1β and IL-8 measurements in proximal small intestinal tissue and colon tissue. **P* < 0.05. CON = control, FVT = fecal virome transfer, CVT = chemostat virome transfer, and CVT-MO = CVT propagated with milk oligosaccharides

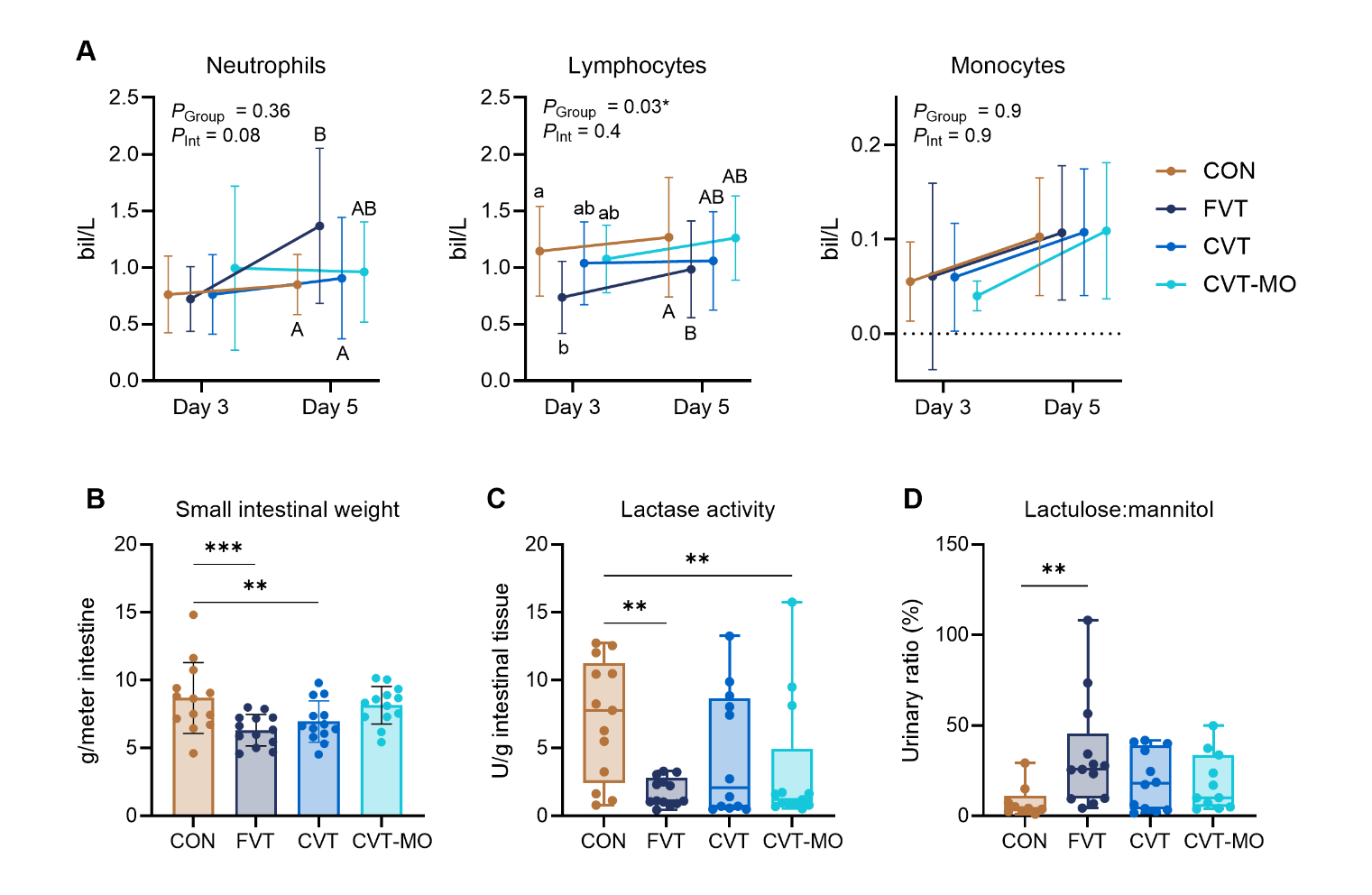

**Supplementary** Figure S6**.** Paraclinical data from *Piglet Experiment 1*. A) Leukocyte counts on days 3 and 5. *P*_Group_ = *P*-value for group effect*, P*_Int_ = *P*-value for group and time interaction. Lines not sharing the same lowercase letter are significantly different (*P* < 0.05). Lines not sharing the same uppercase letter are borderline different (*P* < 0.1). B) Small intestinal weight relative to length. C) Lactase activity in proximal small intestinal homogenates. D) Urinary lactulose to mannitol ratio 3 hours after oral bolus. **P* < 0.05, ***P* < 0.01, ****P* < 0.001. CON = control, FVT = fecal virome transfer, CVT = chemostat virome transfer, and CVT-MO = CVT propagated with milk oligosaccharides.

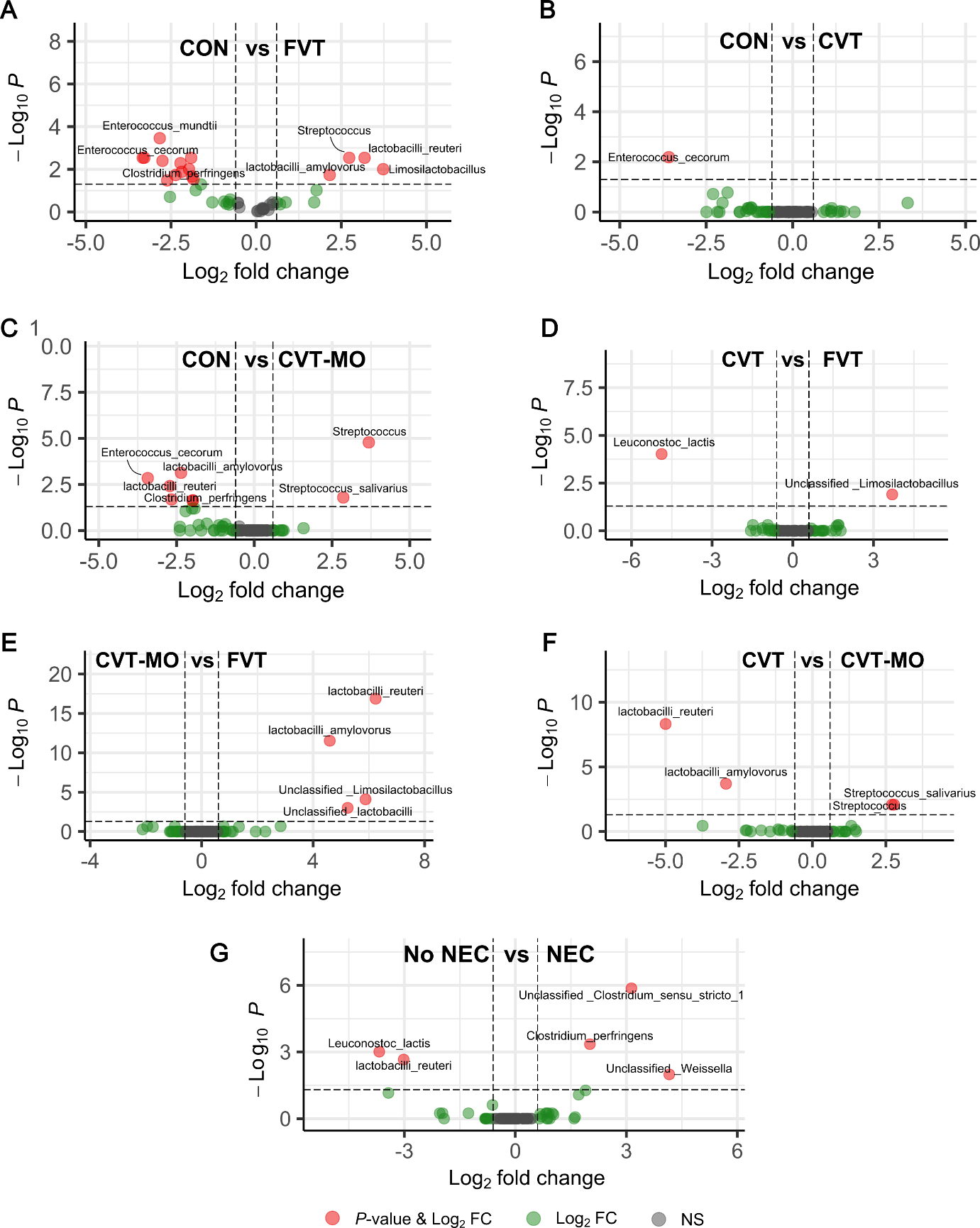

**Supplementary Figure S7:** Vulcano plots showing pairwise DESeq2 enrichment analyses of colon bacterial species based on 16S rRNA gene amplicon sequencing with false discovery rate correction. Horizontal dotted lines show significance level of *P* < 0.05 and vertical lines show ± 0.6 Log2 fold change (FC) cut-off. NS = non-significant. CON = control, FVT = fecal virome transfer, CVT = chemostat virome transfer, and CVT-MO = CVT propagated with milk oligosaccharides (*n* = 10-13/group)

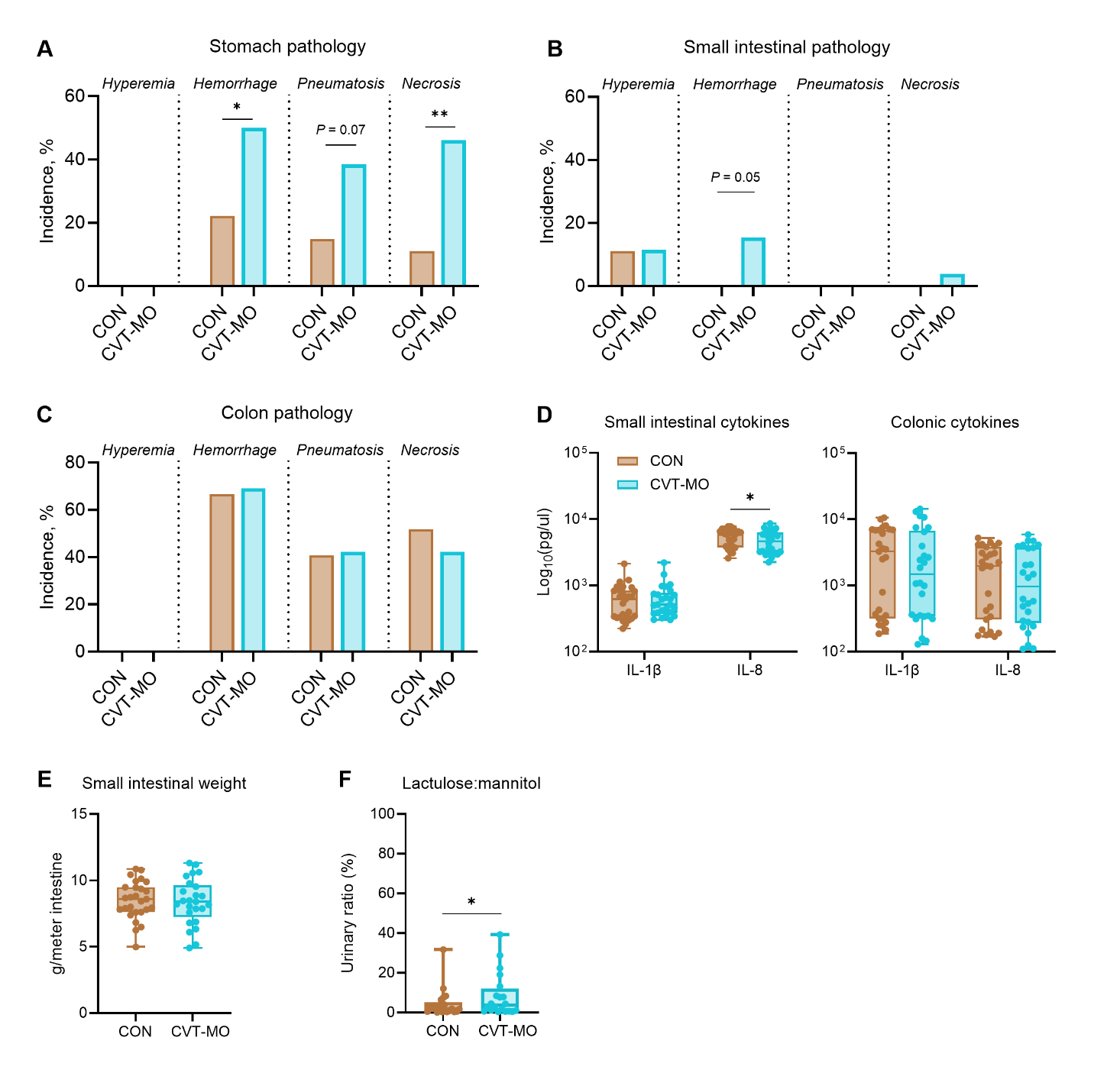

**Supplementary Figure S8**. Supporting pathology data and paraclinical outcomes from *Piglet Experiment 2*. A-C) Incidences of hyperemia, hemorrhage, pneumatosis intestinalis (intramural gas), and necrosis in stomachs (A), small intestines (B), and colons (C). D) Interleukin (IL) 1β and IL-8 measurements in proximal small intestinal tissue and colon tissue. E) Small intestinal weight relative to length. F) Urinary lactulose to mannitol ratio 3 hours after oral bolus. **P* < 0.05, ***P* < 0.01. CON = control, CVT-MO = chemostat virome transfer propagated with milk oligosaccharides (*n* = 26-27).

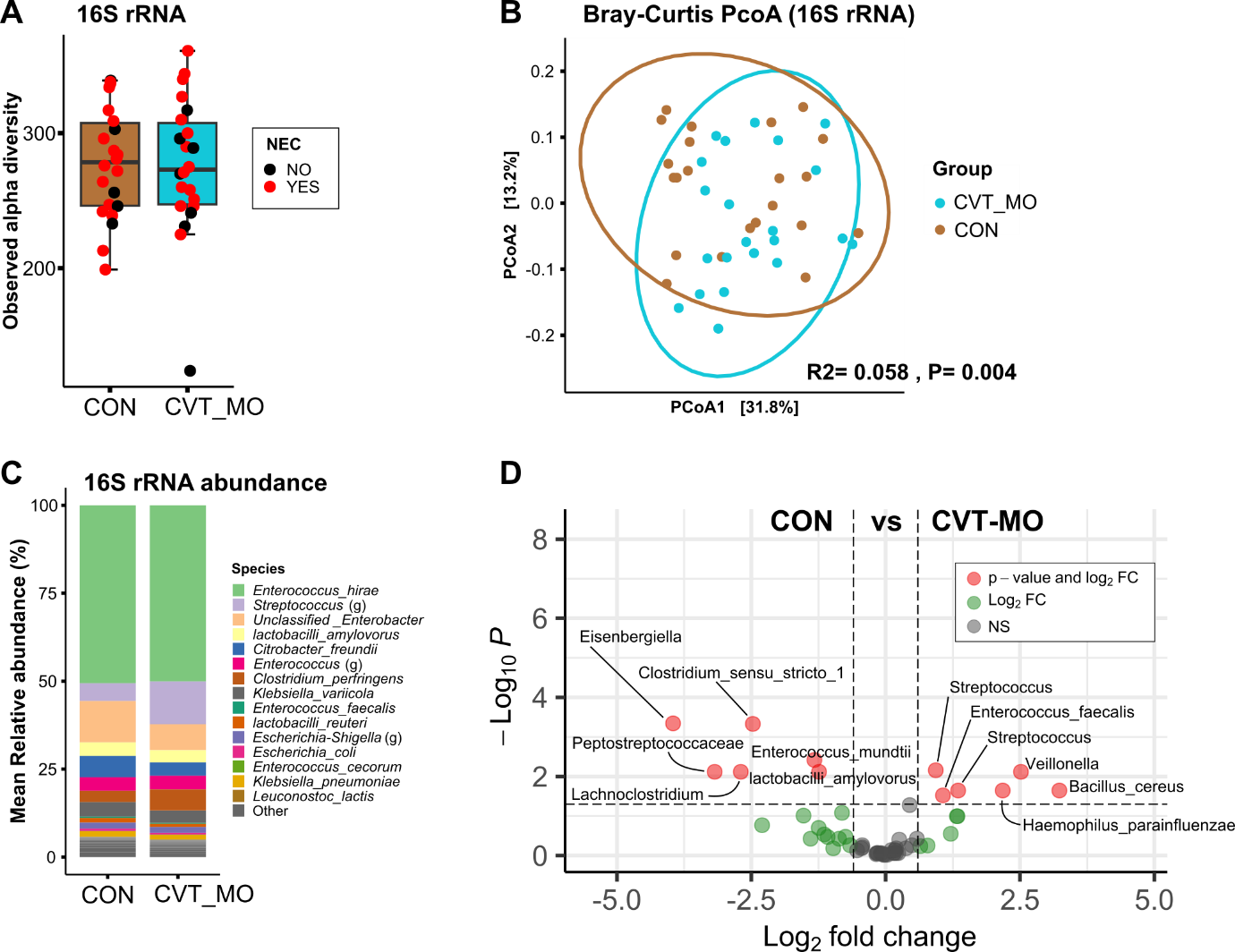

**Supplementary Figure S9.** Bacterial compositions in colons from *Piglet Experiment 2*. A) Number of observed bacterial species as a measure of bacterial alpha diversity between. B) Principal component analysis (PCoA) plot visualizing bacterial beta diversity based on Bray-Curtis dissimilarity metrics. C) Mean relative bacterial abundance summarized at species or genus (g) level. D) Vulcano plot showing pairwise DESeq2 enrichment analyses of colon bacterial species based on 16S rRNA gene amplicon sequencing with FDR correction. Horizontal dotted lines show significance level of *P* < 0.05 and vertical lines show ± 0.6 log2 fold change (FC) cut-off. NS = non-significant. CON = control, CVT-MO = chemostat virome transfer propagated with milk oligosaccharides (*n* = 22).

**Supplementary Tables**

**Supplementary** Table S1. Characteristics of bacteriophages in cocktail used for the THP-1 cell stimulation.

| **Annotation name** | **Genus** | **Morphology** | **Bacterial host** |
| --- | --- | --- | --- |
| vB_Ecoli_M_ECO5 | *Mosigvirus* | Long, contractile tail | *Escherichia coli* ST-2064 |
| vB_Ecoli_M_ECO9 | *Krischvirus* | Long, contractile tail | *Escherichia coli* ST-10 |
| vB_Ecoli_M_ECO12 | *Krischvirus* | Long, contractile tail | *Escherichia coli* ST-10 |
| vB_Ecoli_M_ECO15 | *Tequatrovirus* | Long, contractile tail | *Escherichia coli* ST-10 |
| vB_Ecoli_M_ECO22 | *Krischvirus* | Long, contractile tail | *Escherichia coli* ST-10 |
| vB_Ecoli_P_ECO27 | *Teseptimavirus* | Short, non-contractile tail | *Escherichia coli* ST-2064 |
| vB_Ecoli_P_ECO30 | *Teseptimavirus* | Short, non-contractile tail | *Escherichia coli* ST-2064 |
| vB_Ecloacae_S_ECL3 | *Warvickvirus* | Long, filamentous, non-contractile tail | *Enterobacter cloacae* ST-104 |
| vB_Ecloacae_S_ECL15 | *Warvickvirus* | Long, filamentous, non-contractile tail | *Enterobacter cloacae* ST-104 |
| vB_Ecloacae_S_ECL18 | *Warvickvirus* | Long, filamentous, non-contractile tail | *Enterobacter cloacae* ST-104 |

**Supplementary** Table S2. Primer sequences used for 16S amplicons analysis.

| **Primers** | **Primer Sequence** |
| --- | --- |
| UMI_338Fa | 5’- GTCTCGTGGGCTCGG- NNNNNNNNNNNNNNN - ACWCCTACGGGWGGCAGCAG-3’ |
| UMI_338Fb | 5’- GTCTCGTGGGCTCGG- NNNNNNNNNNNNNNN - GACTCCTACGGGAGGCWGCAG-3’ |
| UMI_27Fa | 5’- GTCTCGTGGGCTCGG- NNNNNNNNNNNNNNN - AGAGTTTGATYMTGGCTYAG-3’ |
| UMI_27Fb | 5’- GTCTCGTGGGCTCGG- NNNNNNNNNNNNNNN - AGGGTTCGATTCTGGCTCAG-3’ |
| UMI_1540R | 5’- GTCTCGTGGGCTCGG- NNNNNNNNNNNNNNN - TACGGYTACCTTGTTACGACT-3’ |
| UMI_1391R | 5’- GTCTCGTGGGCTCGG- NNNNNNNNNNNNNNN -  GACGGGCGGTGTGTRCA-3’ |

**Supplementary** Table S3. Pairwise DESeq2 enrichment analyses from 16S rRNA amplicon sequencing. Ordered from lowest to highest fold change. Only showing species with a *P*-value below 0.05. CON = control, FVT = fecal virome transfer, CVT = chemostat virome transfer, and CVT-MO = CVT propagated with milk oligosaccharides.

| **Log2 fold change** | **Adjusted *P-*value** | **Bacterial classification** |
| --- | --- | --- |
| **FVT vs. CON** | | |
| 3.728883863 | 0.00991103 | Unclassified _Limosilactobacillus |
| 3.182769355 | 0.00290938 | lactobacilli_reuteri |
| 2.731113789 | 0.00290938 | Unclassified_Streptococcus |
| 2.159023194 | 0.01857580 | lactobacilli_amylovorus |
| 1.772713096 | 0.09645004 | Streptococcus_salivarius |
| -1.621376466 | 0.05149558 | Unclassified _Escherichia-Shigella |
| -1.774058787 | 0.09645004 | Unclassified _Veillonella |
| -1.842915916 | 0.02916072 | Unclassified _Citrobacter |
| -1.885002555 | 0.02182549 | Unclassified_Enterobacteriaceae |
| -1.908633626 | 0.00290938 | Escherichia_coli |
| -1.973284833 | 0.00991103 | Clostridium_perfringens |
| -2.188460249 | 0.01318971 | Staphylococcus_aureus |
| -2.221417392 | 0.00518618 | Staphylococcus_epidermidis |
| -2.36555965 | 0.01872935 | Unclassified _Clostridium_sensu_stricto_1 |
| -2.613195648 | 0.03335227 | Leuconostoc_lactis |
| -2.750875308 | 0.00403135 | Enterococcus_faecalis |
| -2.830196362 | 0.00034842 | Enterococcus_mundtii |
| -3.283182112 | 0.00290938 | Unclassified _Rothia |
| -3.328751243 | 0.00290938 | Enterococcus_cecorum |
| **CVT vs. CON** | | |
| -3.58316646 | 0.0064886 | Enterococcus_cecorum |
| **CVT-MO vs. CON** | | |
| 3.682606064 | 0.0000002 | Unclassified_Streptococcus |
| 2.861163841 | 0.0008591 | Streptococcus_salivarius |
| -1.977069137 | 0.0017155 | Clostridium_perfringens |
| -1.983598539 | 0.0020148 | Staphylococcus_epidermidis |
| -2.357327586 | 0.0000158 | lactobacilli_amylovorus |
| -2.665203866 | 0.00131669 | Unclassified _Rothia |
| -2.716106565 | 0.00016455 | lactobacilli_reuteri |
| -3.427194991 | 0.00004604 | Enterococcus_cecorum |
